# Unification of cell-scale metabolic activity with biofilm behavior by integration of advanced flow and reactive-transport modeling and microfluidic experiments

**DOI:** 10.1101/2024.01.25.577134

**Authors:** Jiao Zhao, Mir Pouyan Zarabadi, Derek M. Hall, Sanjeev Dahal, Jesse Greener, Laurence Yang

**Affiliations:** Department of Chemical Engineering, Queen’s University, Kingston, Ontario, Canada, K7L 3N6; Département de chimie, Faculté des sciences et de génie, Université Laval, Québec, Québec; Department of Energy and Mineral Engineering, The Pennsylvania State University

## Abstract

The bacteria *Geobacter sulfurreducens* (GS) is a promising candidate for broad applications involving bioelectrochemical systems (BES), such as environmental bioremediation and energy production. To date, most GS studies have reported biofilm-scale metrics, which fail to capture the interactions between cells and their local environments via the complex metabolism at the cellular level. Moreover, the dominance of studies considering diffusion-only molecular mass transport models within the biofilm has ignored the role of internal advection though the biofilm in flow BES. Among other things, this incomplete picture of anode-adhered GS biofilms has led to missed opportunities in optimizing the operational parameters for BES. To address these gaps, we have modernized a GS genome-scale metabolic model (GEM) and complemented it with local flow and reactive-transport models (FRTM). We tuned certain interactions within the model that were critical to reproducing the experimental results from a pure-culture GS biofilm in a microfluidic bioelectrochemical cell under precisely controlled conditions. The model provided insights into the role of mass transport in determining the spatial availability of nutrient molecules within the biofilm. Thus, we verified that fluid advection within biofilms was significantly more important and complex than previously thought. Coupling these new transport mechanisms to GEM revealed adjustments in intracellular metabolisms based on cellular position within the biofilm. Three findings require immediate dissemination to the BES community: (i) Michaelis-Menten kinetics overestimate acetate conversion in biofilm positions where acetate concentration is high, whereas Coulombic efficiencies should be nearly 10% lower than is assumed by most authors; (ii) unification of the empirically observed flow sensitivity of biofilm-scale kinetic parameters and cell-scale values are finally achieved; and (iii) accounting for advection leads to estimations of diffusion coefficients which are much lower than proposed elsewhere in the literature. In conclusion, in-depth spatiotemporal understanding of mechanisms within GS biofilm across relevant size scales opens the door to new avenues for BES optimization, from fine-scale processes to large-scale applications, including improved techno-economic analyses.

## Introduction

Electroactive biofilms, such as those from *Geobacter sulfurreducens* (GS), are key living catalytic materials in emerging bioelectrochemical systems (BES). Applications of BES include bioremediation of waste streams and environmental sites, such as those containing radioactive elements^1^ as well as inorganic, aromatic, and other hydrocarbon species.^2^ Electroactive biofilms in BES are also intensively studied as energy materials in microbial fuel cells (MFCs)^3^ and for other bioenhanced processes, including desalination, electrolysis, electrosynthesis and methane production.^4^ Pure-culture GS biofilms are recognized as efficient in organic-to-electricity biotransformation.^5^ They are also excellent model biofilms because strain composition is fixed and electron transfer is direct and uncomplicated. In addition, the continuous current production is an accurate proxy for real-time assessment of the average metabolic rate at the biofilm-scale, which enables a range of detailed kinetic studies.^6^ In addition to enabling straightforward analysis of experimental results, GS is an excellent subject in modeling and theoretical studies focusing on fundamental mechanisms.

Among GS modeling approaches, the flow and reactive transport model (FRTM) has been used to create spatiotemporal simulations for predicting GS bioremediation of soluble uranium (U^6+^) to insoluble uranium (U^4+^) minerals present in groundwater via environmental acetate loading.^7^ Monod-type growth kinetics were used to represent various multi-enzyme cycles as a general black box in the FRTM, which linked the reduction rates of extracellular electron accepters (e.g., Fe^3+^ and U^6+^) to the acetate oxidation rates through universally applied electron-transfer ratios (ETR) and biomass yields (Y_bio/ace_). However, the FRTM model did not account for the reality that spatiotemporal environmental conditions may influence the metabolism of the GS bacteria at different locations within the biofilm. Some work has partially addressed this gap with genome-scale metabolic modeling (GEM), which integrated the GS intracellular metabolic network with the local extracellular environment for bioremediation applications.^8^ Their GEM used a metabolic flux balance analysis (FBA) to calculate the optimal reduction rates and biomass production rates from the rates at which acetate oxidation proceeded through intracellular metabolism. Hence, the reaction stoichiometry, yield, and transfer coefficients in that study could vary to reflect the ability of GS cells to respond to their local environment. However the model did not capture the entire 3-dimensional porous structure of a real GS biofilm because the spatial resolution was at the decimeter scale or larger,^8^ whereas thicknesses of metabolically active biofilms are less than a fraction of a millimeter.^9^ Moreover, GEM generally represents the cellular metabolic response to its immediate extracellular environment. To this end, we refer to the “cell-scale” as the scale at which cells interact with their local environment. To reduce this gap in scale, a subsequent example from the literature coupled GEM to FRTM (GEM-FRTM) through the soil pore spaces.^10^ However, this simulation was computationally intensive due to the small soil pore dimensions, so the authors had to increase all reaction rates in their model by a factor of 10^3^ to speed up the coupling between biotic and abiotic processes. Again, the model did not have sufficient spatial resolution to consider the GS within the biofilm 3D matrix and therefore could not account for variations in metabolic activities within the biofilm.

A more useful model for BES should consider cell accumulation into electrode-adhered biofilms with thicknesses at an appropriate length scale that can account for internal concentration gradients, effective diffusivity, and advection through the biofilm. It is known that electrode-adhered GS can develop metabolically active biofilms with thicknesses greater than 50 µm^9^ and average porosities greater than 0.45.^11^ Reduced diffusional mass transport of acetate ions (CH_3_COO^-^), the major substrate for GS, in biofilms with thickness greater than 45 µm has been associated with limitations to electrochemical activity.^12^ This observation is corroborated by another study that showed a stratification of fluorescent tracers between accessible regions near the biofilm/liquid interface and inaccessible regions deeper in the GS biofilm.^13^ One can understand such limitations in diffusion as being caused by acetate confinement in pores and increased molecular adsorption and simultaneous consumption in dense bacterial clusters.^14, 15^ The diffusion of negatively charged acetate ions may also be influenced by electrostatic interactions in these confined pore spaces within the GS biofilm.^16^ Given that all mass transport inside a biofilm is traditionally attributed to diffusion in the BES literature^11^ and that GS biofilm properties vary widely based on age and culture conditions,^17^ it is no surprise that missing contributions to other mass transport mechanisms has resulted in such a wide range of diffusivities being reported in the literature (0.7–9.0 mm^2^ h^-1^).^11, 15^

We propose microfluidic BES as the best experimental platform for probing advection through the GS biofilm due to the limited ability for flow redirection around it. This couples well with several other advantages of microfluidic experimental platforms in studying biofilms and offers benefits from real-time measurements while exercising precise control over experimental conditions (e.g., hydrodynamics and applied concentrations).^18–20^ As such, microfluidic bioelectrochemical devices have led to a better fundamental understanding of the coupled physical and biological processes in BES,^21^ especially when combined with numerical models and theory that can account for kinetic and mass transport in and around the biofilm.^18, 22–24^

While the spatial resolution of most microfluidic-scale models is adequate to discern the biofilm structure, such models used to study reaction kinetics mostly use empirical Michaelis-Menten (M-M) kinetics, or M-M-derived kinetics, to determine metabolic activities. However, the lack of direct coupling between hydrodynamics, mass transport, and intracellular kinetics is hypothesized to cause an incorrect estimation of the metabolic response to a changing extracellular environment at the cell-scale. Challenges also arise in relating mechanisms at the biofilm-scale to those at play at the cell-scale, especially when accounting for advective flow through the biofilm. Specifically, relating to, (1) how the parameters in empirical M-M kinetics at the biofilm-scale relate to reaction kinetics and stoichiometry at the cell-scale and (2) the role of internal advection on GS biofilm performance.

To address these open questions, we modeled a GS biofilm that developed to a thickness of 80 µm in a microfluidic device to study the effect of an acetate nutrient flow on the overall biofilm performance for generation of electric current. Based on the experimental setup, we developed a full microfluidic model that couples biological phenomena at the cell-scale to the local extracellular mass transport phenomena, both of which are difficult to study experimentally. We found that accounting for advection within the biofilm was critical in reproducing the results from microfluidic experiments. In doing so, we discovered that the acetate diffusion coefficients for our GS biofilm are as low as 0.05–0.30 mm^2^ h^-1^, depending on the flow and acetate conditions. This is much lower than previously reported (0.7–9.0 mm^2^ h^-1^)^11, 15^ for GS or general biofilms under flow conditions, in which advection inside the GS biofilms was ignored. The GEM-FTRM approach for the GS model also predicts that at high acetate concentrations, the cellular electron production per acetate molecule is only 7.33, which is almost 10% lower than is typically assumed based on the chemical stoichiometry of acetate oxidation. Lastly, previous experimentally observed reductions to the M-M constant 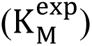 when GS biofilms were subjected to increased flow rates^23, 25^, which were successfully predicted by extrapolating from the cell-scale M-M constant 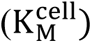 to the biofilm-scale 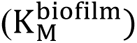 using GEM-FRTM simulations.

## Results and discussion

### Improvement of GEM for GS

To use GS GEM as a reliable tool for the study of the electron and mass flow between the intracellular metabolic network and its surrounding extracellular environment, the first goal was to modernize the previously updated GEM for GS (GS_v2),^26^ which was based on the original GS_v1.^27^ This was accomplished by converting the GS_v2 into human-readable BiGG form,^28^ adding the charge balance for all intracellular metabolites, removing orphan intracellular metabolites (which are only consumed but not produced in the GEM) as well as their associated reactions, and correcting the names for erroneous gene–protein-reaction associations. The modernized GS GEM structure is outlined in GS_v3.xlsx in Electronic Supplementary Information (ESI).

We conducted quality control of our modernized GS GEM version (GS_v3) using the open-source Python-based metabolic model testing tool (MOMOTE).^29^ The quality report is shown in Table S1 in the ESI. MOMOTE reported 100% consistency in stoichiometry, mass balance, charge balance and metabolite connectivity. MOMOTE identified 5.62% (15 reactions) of the reactions that may carry unbounded metabolic fluxes, which are involved in thermodynamically infeasible loops. The Loopless-FBA approach^30^ by mixed-integer linear programming (MILP) algorithm was used to eliminate these thermodynamically infeasible loops found in FBA. The results were compared with those obtained by the AllowLoops-FBA approach by linear programming (LP) algorithm. It was found that the 15 unbounded intracellular metabolic fluxes had no effect on the GS phenotypes represented by all extracellular metabolic fluxes.

Thus, our modernized GEM reliably traces the electron flow through the intracellular redox metabolism and relates it correctly to the extracellular current generation and biomass yield. In achieving this improvement, we discovered that the commonly assumed electron transfer ratio (ETR; the stoichiometry governing the number of electrons produced per acetate molecule oxidized) is not always 8 electrons per molecule, as commonly assumed, but continuously varies with acetate concentration. As discussed below, this has major implications for the BES research community due to its impact on expected Coulombic efficiency. It also opens the way for optimizing our GEM-FRTM.

The new GEM of a GS cell (GS_v3) consists of 466 metabolites, 538 reactions, 889 genes and 367 reactions with gene–protein-reaction association. The improved GS GEM is provided in the ESI in SBML, EXCEL and MATLAB MAT format.

### Reduced computational costs for GEM-FRTM

The GEM-FRTM at a fine spatiotemporal scale is computationally intensive because the GEM must iteratively call linear programming (AllowLoops-FBA) or mixed-integer linear programming (Loopless-FBA) once per iteration more than a million times during a simulation for all discrete points and time steps. In our study, the GEM-FRTM simulation by AllowLoops-FBA took approximately 8.3 hours of CPU time (Intel Core i7-10875H @ 2.30 GHz) to describe approximately 2.7 minutes of the process occurring in the microfluidic system, which makes long-term large-scale model simulations impractical due to the unacceptable computational cost. The Loopless-FBA further increases the computational cost by 10-fold. To solve this problem, two polynomial look-up functions were generated, which could avoid LP or MILP computations. This was accomplished by making practical use of the previously noted concentration-dependent GEM-derived electron transfer ratio (ETR) and the biomass yield (Y_bio/ace_; g of biomass dry cell weight produced per mmol of acetate oxidized) based on 10^4^ evenly distributed acetate concentrations between 0 and 10 mM using an n^th^ degree polynomial regression (Figure 1A and 1B by the AllowLoops-FBA). The polynomial function, when coupled to the FRTM, was then called at each time step and each grid point to compute ETR and Y_bio/ace_ in the local extracellular environment. Figure 1C indicates that such an indirect GEM coupling method (iGEM) can exactly duplicate the full GEM coupling method, and the invested computational costs are similar to that of the more simplistic but faster Monod-type M-M coupling method that treats intracellular metabolism as a “black box” (see M-M coupling in methods section). In other words, iGEM achieves the best aspects of GEM (accuracy) and M-M modeling (speed). As an example, using iGEM, 8.3 hours of CPU time simulated up to 81 minutes of the real process, which is approximately 30 times faster than the direct GEM method (Figure 1D).

**Figure 1.**
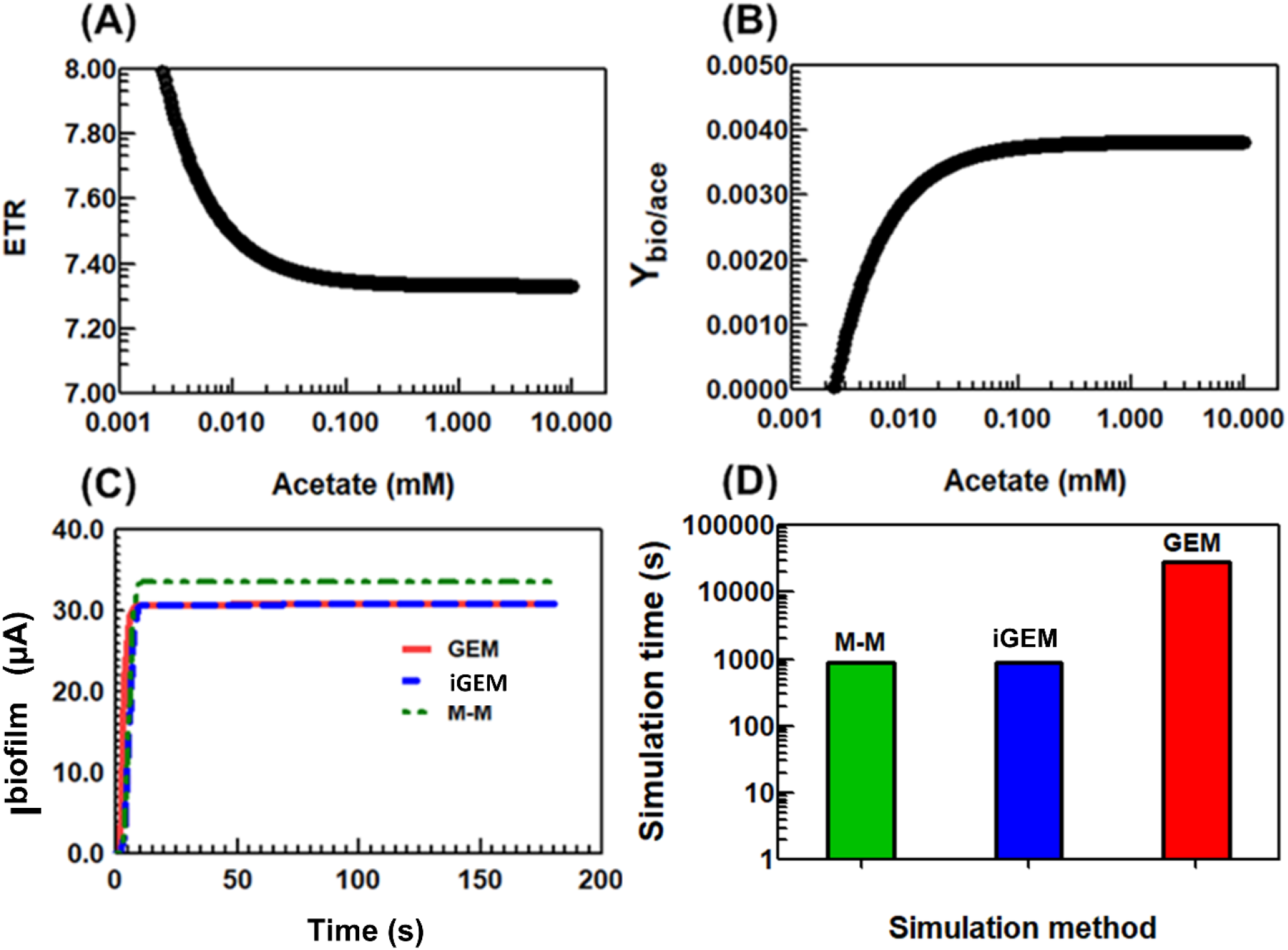
Assessment of model performance. Indirect GEM (iGEM): an algorithm that indirectly couples GS GEM to mass transport equations by GEM-predicted metabolic activities as a function of local acetate concentrations. (A) Semi-log regression of GEM-predicted electron transfer ratio (ETR, number of electrons transferred to current per oxidized acetate molecule) as a function of local acetate concentration. (B) Semi-log regression of GEM-predicted biomass yield (Y_bio/ace_, g dry-cell-weight biomass produced per mmol of acetate oxidized) as a function of local acetate concentration. See the ESI for 6^th^-degree polynomial fitting functions used in A and B. (C) Simulated I^biofilm^ from direct Michaelis-Menten (M-M, green), indirect GEM coupling algorithm (iGEM, blue), and GEM coupling algorithm (GEM, red) under 8.0 mL h^-1^ flow rate of 10 mM acetate. (D) Computational cost for M-M, GEM and iGEM coupling algorithms with same color-coding as in (C).

For comparison, the pre-calculated Loopless-FBA predictions under 10^4^ possible concentrations of acetate are shown in the ESI (Figure S1). Under test microfluidics conditions, the thermodynamically infeasible loops found in the GS GEM had no effect on ETR and Y_bio/ace_, however, they should be eliminated for energy balance purposes to avoid the risk of mass and charge inequality for some GEM reactions in a large system. Unfortunately, the Loopless FBA approach is computationally expensive due to the MILP algorithm, which makes the coupling of GEM to FRTM impractical. The iGEM method proposed in this study provides an effective solution to addressing such computational expenses in case the Loopless-FBA is mandatory for the GEM. To exploit the iGEM to study this and other phenomena within the GS biofilm, we propose the use of parallel computing at discrete points in biofilms to further speed up biofilm simulations as the next step towards an iterative model-based on-site experimental optimization.

We note with interest that the changes to ETR are expected based on the local acetate concentration (Figure 1A). Moreover, Figure 1C shows that the simulated electric current from the entire GS biofilm (I^biofilm^, µA) predicted from the M-M method was always approximately 10% higher than that from the GEM and iGEM methods under high-acetate and high-flow-rate conditions. The mechanism behind this phenomenon will be explored in the discussion section.

### Parameter tuning using experimental results

With iGEM coupled to FRTM (iGEM-FRTM) in hand, we sought to iteratively tune the model parameters starting with experimentally determined values found in the literature (Table 1) and reasonable estimates (ESI). The simulation parameters included the local mass transport values around the GS cell surfaces (i.e., the cell-scale), namely, diffusion (D, mm^2^ h^-1^), permeability (k, mm^2^), and porosity (φ, unitless), and the biological kinetic parameters at a cell-scale, namely, the maximum cellular reaction rate at infinite concentration (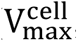, mmol g^-1^ h^-1^), the cellular half-maximum rate concentration (otherwise known as the M-M constant, 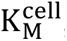, mmol mm^-3^), and the initial acetate-accessible catalytic biomass concentration (BMC, g mm^-3^). It is important to note that the kinetic parameter 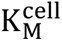 included in the iGEM was obtained from a well-mixed continuous reactor for planktonic GS cells,^31^ and thus, it can be considered a fundamental cell-scale constant with lower measurement uncertainty.

**Table 1.**
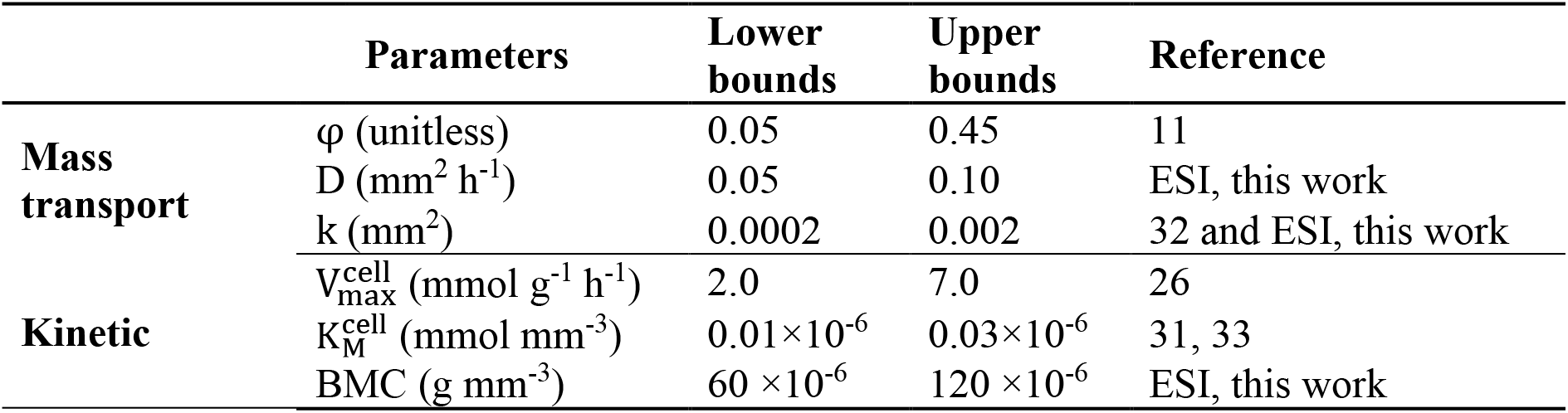
Range of key kinetic and mass transport parameters values in the microfluidic iGEM-FRTM model.

As a starting point, we assumed that these parameter values in a biofilm were constant and independent of space, time, and experimental conditions. In addition, we assumed that the mass transport inside the GS biofilm is only driven by diffusion without advection, as was assumed in a previous study of a GS biofilm using a flow-NMR reactor.^11^ The complementary experiments in this work enable comparison with the model outputs, which, as we show later, were used to test these assumptions.

Our simulated current profiles, integrated across the entire biofilm (I^biofilm^), indicate flow-insensitivity (Figure S2 in ESI) under low flow rates between 0.4 and 2.0 mL h^-1^ for high acetate concentrations when a wide range of D from 0.05 to 5.0 mm^2^ h^-1^ was used. This is contrary to what has been observed experimentally at these flow rates (red dots in Figure S2), where the total experimental current (I^exp^) was found to increase with flow rates at high acetate levels ranging from 7 mM to 10 mM.

However, when we introduced flow to the biofilm and decreased D appropriately to account for the parallel mass transport mechanism via advection through the GS biofilm, the iGEM-FRTM-derived I^biofilm^ was able to reproduce the sensitivity of the measured I^exp^ to the lower flow rates (from 0.4 to 2.0 mL h^-1^) at higher acetate concentrations (from 7.0 to 10.0 mM) (Figure 2). This is accomplished by introducing a permeability, k which is used by the Navier-Stokes-Brinkman equation to represent the ability of the biofilm to support advective mass transport while decreasing D to approximately 0.05 mm^2^ h^-1^.^32^ In the absence of reported k values for GS biofilms, we used the maximum and minimum k values found for general biofilms as the upper and lower bounds (see ESI). The red shaded areas in Figure 2 show the I^biofilm^ solution space that was obtained using k values within these limits. This solution space indicates that the simulated I^biofilm^ was lower than I^exp^ under low-acetate (0.3 mM) and low-flow conditions (0.4 ∼ 2.0 mL h^-1^) (Figure 2A), even if the upper bounds of φ, D and k were used (Table 1). However, the I^biofilm^ was higher than I^exp^ under high acetate concentrations (10 mM) (Figure 2D) even if the lower bounds of φ, D and k were used. Moreover, as acetate concentration increased, the permeability k had to be decreased to achieve a good fit between simulations and measurements. These facts suggest that the GS biofilm internal architecture (e.g., φ) and mass transport parameters (e.g., D and k) may change with experimental conditions. Thus, the constant values we used for parameter tuning may not correctly describe the biofilm performance under different experimental conditions.

**Figure 2.**
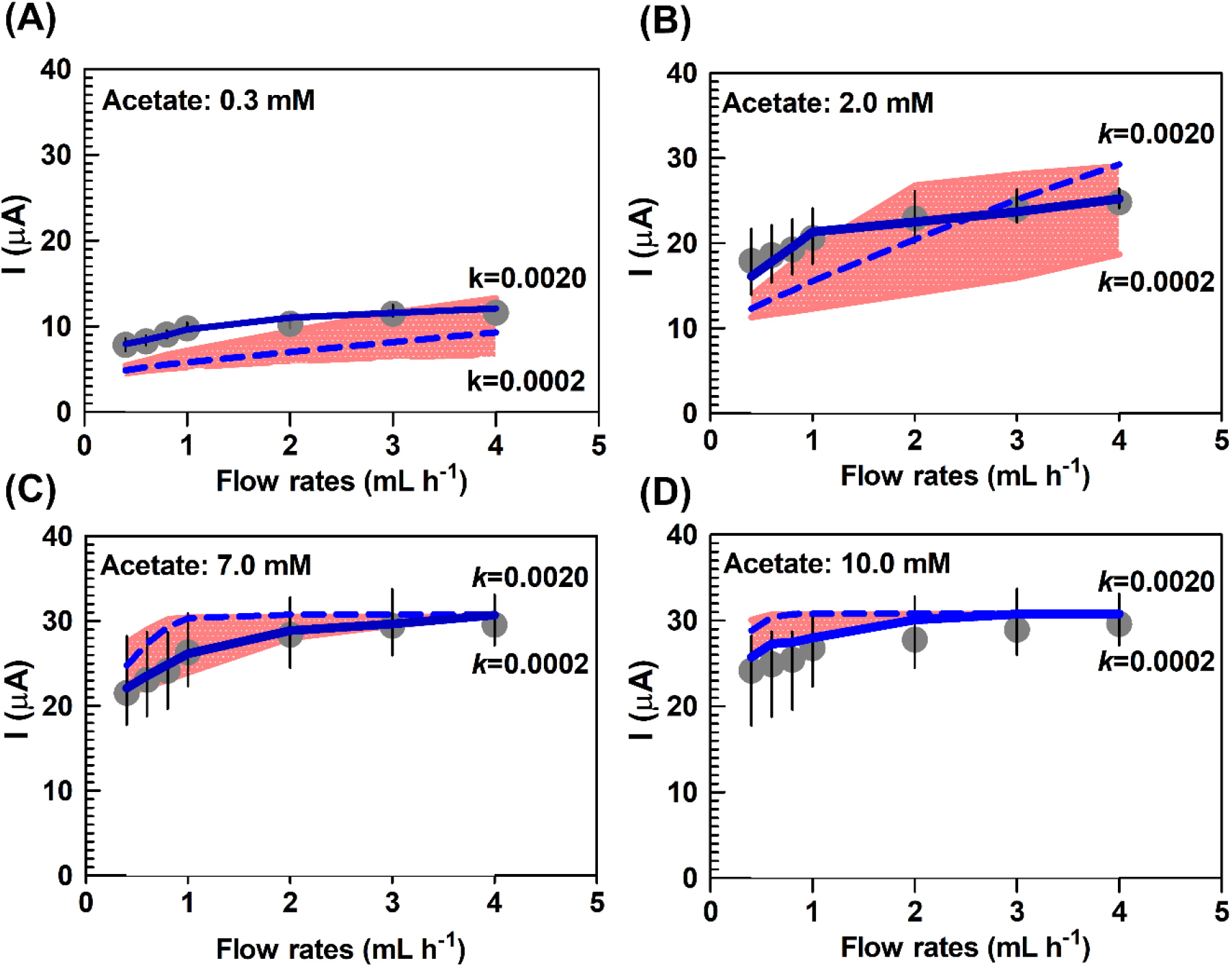
Comparison of currents (I) obtained from simulations and experiments at different flow rates using acetate concentrations of (A) 0.3, (B) 2, (C) 7 and (D) 10 mM. Measured current (I^exp^, gray circles) includes error bands, which were obtained from the standard deviation from three repeat measurements. I^biofilm^ were simulated using either iGEM-FRTM with advection rate set at 10% of the inflow rate (blue dashed lines) or with dynamically optimized changing mass transport parameters φ, D and k due to adjustments of the biofilm matrix to flow rates and acetate concentrations (solid blue line). A range of simulated currents are also shown using permeability values in the range of k=0.0002 mm^2^ to k=0.002 mm^2^ with constant porosity (φ=0.45) and diffusivity (0.05 mm^2^ h^-1^) for all flow rates and acetate concentrations (red band).

Previous studies have reported biofilm structural and mass transport changes with flow,^34^ pH,^35^ and nutrition levels,^36^ including those which support the idea of parameters that are dependent on experimental conditions.^36,37^ It has been observed that an increase in the fluid flow rate and nutrient concentration and a decrease in pH can cause biofilms to deform in complex ways,^38^ leading to a decrease in biofilm φ, D and k.^34, 39^ It should be noted that biofilm mechanical properties are complex^37, 40^ and they will dictate the degree to which the biofilm can respond to applied flow forces, especially in microfluidic conditions.^41, 42^

Based on experimental observations,^34, 36, 39^ we removed the upper bound limits for φ, k and D to accommodate conditions under which biofilms have been shown to become “loose”, an architecture that promotes mass transport towards the accessible biomass catalytic sites. We re-tuned the parameters, this time allowing the fitting protocol to produce dynamic changes to φ, D and k based on the reliable experimental conditions, which were set using a high-precision microfluidic setup. We obtained the best match between I^biofilm^ and I^exp^ (Figure 2) using the profiles of φ, D and k shown in Figure S3 (see ESI).

The mass transport profiles (Figure S3) for low acetate concentration (0.3 mM) suggest that lower flow rates are favorable for the formation of a more loose and more permeable biofilm, which promotes mass transport. Conversly, under the same low acetate concentration, higher flow rates such as 4 mL h^-1^ will compact the biofilm, thus impeding mass transport. However, the mass transport profiles for high acetate concentration (10 mM) are stable at their lower bounds, suggesting that the biofilm tends to densify under higher concentrations. This is likely a biological response to adequate nutrient concentrations for metabolic activity, without the need to saccrafice the dense protective biofilm strucutre, whereas at low concentrations, loss of denity may be an acceptable trade-off to improve mass transport of limited nutrients. Another explaination of the changing biofilm density as a function of flow and concentration may be related to acidification. As certain of our authors have shown before, high acetate levels can result in strong acidification of the GS EAB, which cannot be modulated with flow rate.^25^ Other studies have shown that the acidic environment can counteract charge repulsions in the biofilm, causing the biofilm to become more compact.^35^ Thus, acidic conditions at high acetate concentration may reduce the negative charge requlsions (e.g., between GS outer membrane cytochomes) in the GS biofilm, promoting formation of the compact GS biofilm with reduced mass transport efficiency. We decided to trace the fate of protons in a future iGEM-FRTM version so that these pH-dependant effects can be tracked.

To further understand how acetate concentration along with mass transport influences other factors such as the active biomass concentration, a global sensitivity analysis among model parameters was investigated and presented in the next section.

### Assessment of the individual and integrated impact of kinetic and mass transport parameters on current production

Given the range of values for the model parameters found in the literature and reasonable estimation (Table 1 and ESI), we used the Morris method to quantify the critical role of these parameters applied to local environments through a few iGEM-FRTM evaluations.^43^ This task was accomplished by assessing the effect of both kinetic and mass transport factors on the model output, i.e., I^bioflim^. Figure 3 shows the Morris assessment of the six key input factors under different operating conditions, with increasing values along the x-axis and the y-axis indicative of higher impact of the individual factors (µ*) and higher interaction effects of the individual factors (σ), respectively.

**Figure 3.**
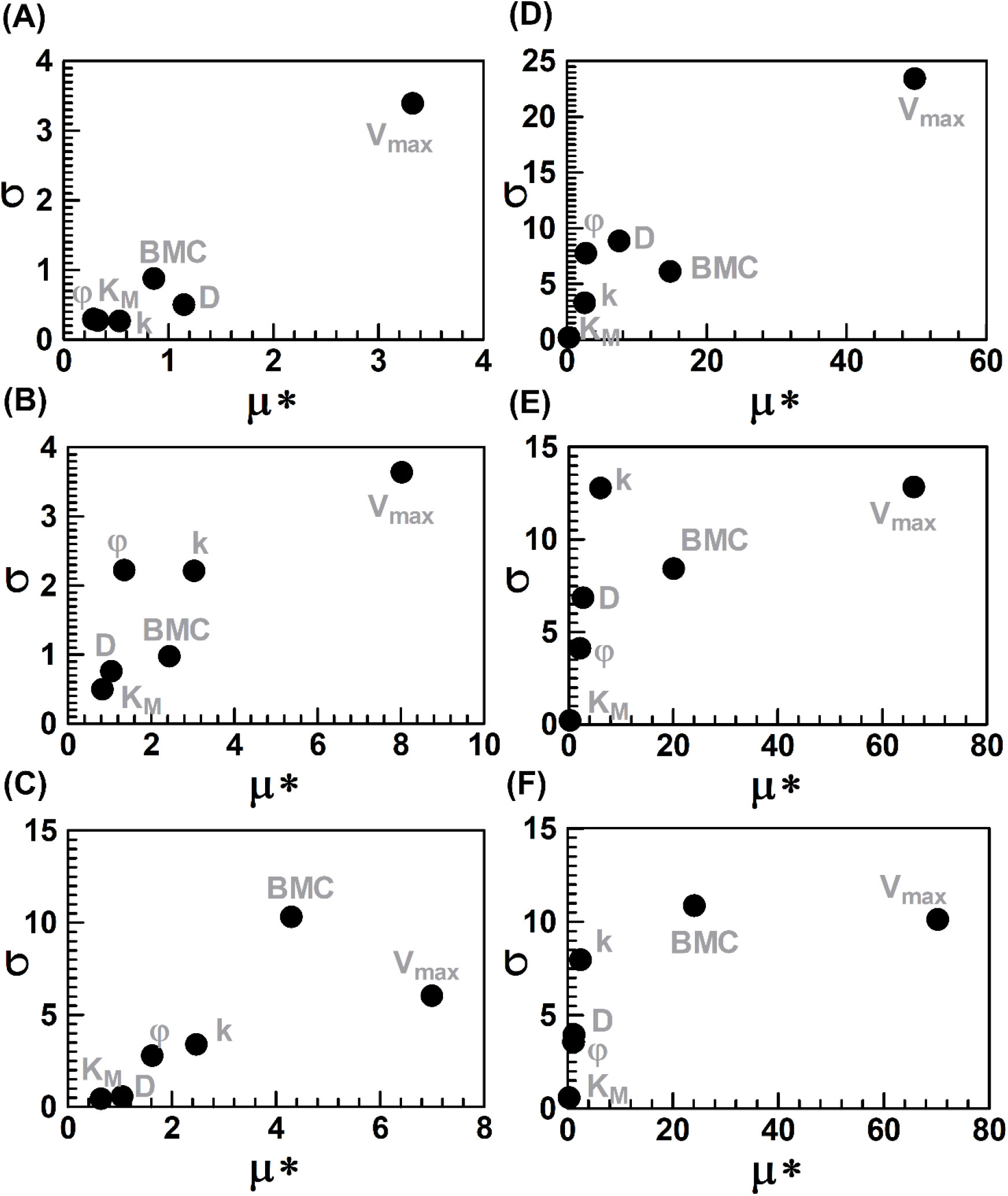
Assessment of key model parameters for local environments. The Morris method shows the individual effect (µ*) and integrated effect (σ) of mass transport (D, k, and φ) and biological kinetic parameters (V_max_, K_M_, and catalytic BMC) on I^biofilm^ in the microfluidics device. (A) 0.4 mL h^-1^ acetate (0.3 mM), (B) 2.0 mL h^-1^ acetate (0.3 mM), (C) 4.0 mL h^-1^ acetate (0.3 mM), (D) 0.4 mL h^-1^ acetate (10.0 mM), (E) 2.0 mL h^-1^ acetate (10.0 mM), and (F) 4.0 mL h^-1^ acetate (10.0 mM).

Values of µ* and σ indicate that except for the values of 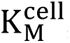, which were obtained with relatively high precision under well-controlled conditions for suspended GS cells,^31^ each of the parameters evaluated had a strong individual or integrated impact on I^biofilm^ within a given numerical range examined (Table 1). The Morris method confirmed that three mass transport parameters (D, k, and φ) and two kinetic parameters (BMC, 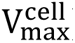) have significant effect on the simulated I^biofilm^, and thus their impact should be closely explored.

First, the individual effect of k is generally more important to I^bioflim^ at high flow rates under low acetate conditions (as observed in the previous section), but the effect of D is more significant under low flow rates (Figure 3A to Figure 3C). These results support our parameter tuning (Figure 2) in that the advection can be more effective in compact biofilms under higher flow rates by supplying acetate deep into biofilm, whereas the diffusion effect is more important in loose biofilms (low acetate concentration) when advective mass transport is insignificant (under low flow rates).

Second, the individual contribution of k to I^bioflim^ (µ*) does not increase with flow rate under high acetate concentrations (Figure 3D to Figure 3F). However, the integrated effect (σ) of both k and BMC becomes more significant as flow rates increase, with σ of BMC reaching the highest value at 4.0 mL h^-1^ of 10 mM acetate. The fact that BMC can work cooperatively with k to increase I^bioflim^ under high acetate concentration suggests that an improved mass transport by advection may increase BMC due to deeper penetration in a compact biofilm, because there may exist regions of the biofilm that were previously starved of acetate which become activated with flow.

As biofilm size can increase from microfluidic devices to macroscale biofilm bioreactors, heterogeneity is likely to lead to spatial variations in k and φ. Therefore, critical assessments of these structural parameters will be helpful for improving the iGEM-FRTM accuracy for application-scale (larger) biofilms. An extended version of the iGEM-FRTM was developed to deal with such heterogeneities as a function of the x- and y-coordinates in 2D (Figure S4). The extended iGEM-FRTM simulations suggest that the biofilm spatial structures may have a significant effect on the electricity generation (Figure S4), fluid dynamics and concentration gradients (Figure S5). Therefore, they cannot be simplified as spatial-independent parameters in large-scale biofilm bioreactors.

### Discussion of new insights gained from iGEM-FRTM

Using the optimized model and model parameters, we evaluated three important questions from first principles: (1) Is the common assumption of 8 electrons per acetate molecule justified? (2) Can the bottom-up iGEM-FRTM starting at a cell-scale move to the biofilm-scale and reproduce the experimental observations by integration? (3) What is the nature of the mass transport and kinetic parameters in iGEM-FRTM?

#### (1) Coulombic efficiency is not constant for different concentrations

Returning to the observed ETR change (Figure 1), when the acetate supply is increased, ETR can drop from 8 to 7.33, which is a reduction of nearly 10%. This corresponds inversely with the production of GS biofilm biomass yield, which is promoted at high acetate concentration. Moreover, the I^biofilm^ predicted from the M-M method was always approximately 10% higher than that from the GEM and iGEM methods under high-acetate and high-flow-rate conditions, suggesting that the M-M method neglecting Y_bio/ace_ increase and ETR reductions at high acetate concentrations may be the cause of deviation.

The equation for chemical oxidation of acetate is given in Eq. 1:

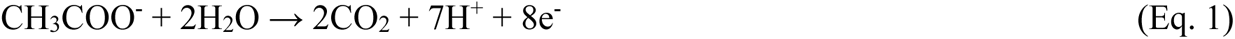

While the constant stoichiometry in Eq. 1 is assumed for much BES work, including as a benchmark for Coulombic efficiency calculations, this may not represent the stoichiometry of biotic oxidation because electrons may be diverted to other biological processes (e.g., biomolecular productions and cell division). A major advantage of the GEM approach is its ability to predict adjustments in the cellular metabolic network in response to extracellular environments, quantify the resulting cell phenotype alterations from 56 exchange fluxes between the intracellular metabolism and the surrounding environment (see GS_v3.xlsx in ESI), and then give better benchmarks for electron production. Moreover, iGEM coupling to the FRTM leverages this capability over the entire biofilm interior. Figure 4 illustrates the metabolic response to acetate availability in 3 different locations within the biofilm. The simulation indicates that in response to the admitted 10 mM acetate solution flowing at 0.2 mL h^-1^ (Figure 4A), orders of magnitude variation in acetate concentration were found throughout the biofilm at steady state, even between some points that are relatively close together. For example, Figure 4A shows the locations where the acetate concentrations were 5 mM, 0.05 mM and 0.002 mM at model sampling points ①, ② and ③, respectively. Figure 4B shows the metabolic flux redistribution estimated by GEM in response to these three acetate concentrations, where 10 non-zero exchange fluxes are used to calculate the stoichiometry for the microbial oxidation of acetate (Figure 4C and Table 2). At low acetate concentration in downstream positions near the electrode surface (0.002 mM), the GS cells approached the theoretical maximum electron transfer ratio (ETR) at 7.99. At these concentrations, most of the acetate was used to meet the demands of cellular energy (ATP) maintenance, with little acetate left for synthesis of new biomass (biomass yield of 0.00005). Because ATP generation via the TCA cycle is directly coupled to electron transfer (Figure 4B), the theoretical maximal electron transfer was reached. The simulated metabolic fluxes (mmol g^-1^ h^-1^) confirm that the relative ATP yield (i.e., V_ATP_/V_ace_ = [mmol ATP g^-1^ h^-1^]/[mmol ace g^-1^ h^-1^] = [mmol ATP]/[mmol ace], representing mmol of ATP produced per mmol of acetate oxidized) increased from 0.44 to 0.50 as the surrounding acetate concentration decreased from 5 mM to 0.002 mM. At 0.002 mM, GS directed 99.9% of the acetate consumed to the TCA cycle via acetate CoA-transferase, compared to 94.1% to TCA at 5 mM acetate. At a higher acetate concentration (5 mM), more acetate was available to increase the rates of both biomass synthesis and ATP generation for electron transfer (Figure 4B). The biomass yield increased to 0.0038 as a result. However, because biomass synthesis requires balancing of energy and anabolic metabolism, more acetate was diverted away from the TCA cycle, thereby reducing ETR from 8 electrons per acetate molecule to 7.33, i.e., a Coulombic efficiency of 92%.

**Figure 4.**
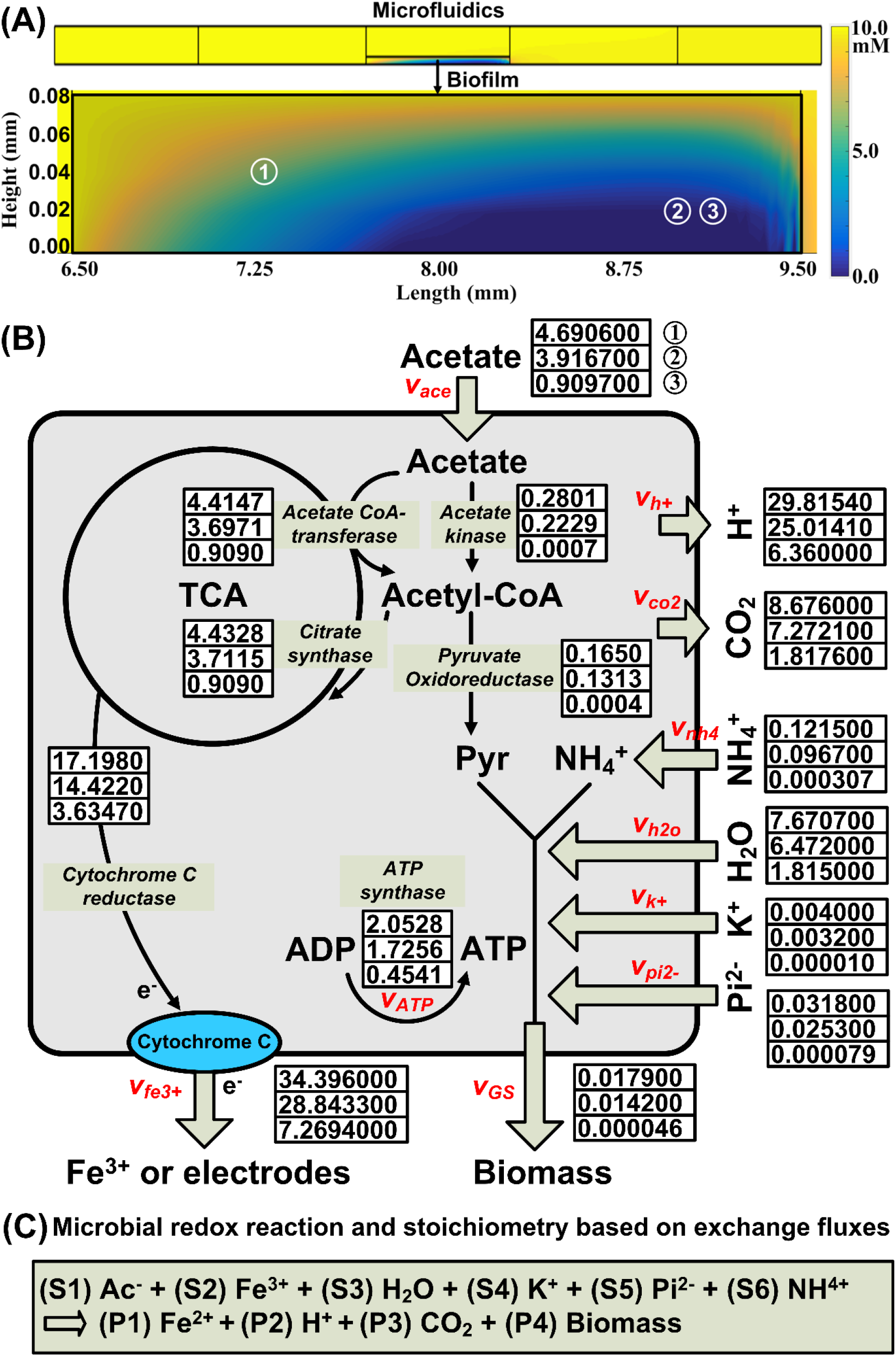
Effect of acetate concentration in local environments on the intracellular metabolic flux redistribution, cellular substrate uptake and production fluxes (exchange fluxes) and the general redox reaction stoichiometry. (A) FRTM of a microfluidic channel filled with acetate (10 mM) flowing at 0.2 mL h^-1^ (top) and acetate concentration distribution over the zoomed biofilm domain (bottom); ①, ② and ③ are sampling points for local environments to calculate metabolic fluxes; (B) GS metabolic flux redistribution at sampling points ①, ② and ③ representing different acetate levels in the biofilm; metabolic fluxes are given in units of mmol g^-1^ dry cell weight h^-1^, except for the GS growth rate (V_GS_) in unit of h^-1^. (C) General redox reaction stoichiometry for substrate consumption (S1-S6) and products (P1-P4); stoichiometric constants are estimated from exchange fluxes (V with substrate or product names in subscript letters; units: mmol g^-1^ dry cell weight h^-1^) through intracellular metabolism, where S1=V_ace_/V_ace_, S2=V_fe3+_/V_ace_, S3=V_h2o_/V_ace_, S4=V_k+_/V_ace_, S5=V_pi2-_/V_ace_, S6=V_nh4_/V_ace_, P1=V_fe2+_/V_ace_, P2=V_h+_/V_ace_, P3=V_co2_/V_ace_ and P4=V_GS_/V_ace_. For simplicity without loss of generality, Fe^3+^ is chosen to be an extracellular electron acceptor, with the same role as outer-membrane cytochromes in the biofilm matrix, to carry electrons towards the anode to generate current.

**Table 2.**
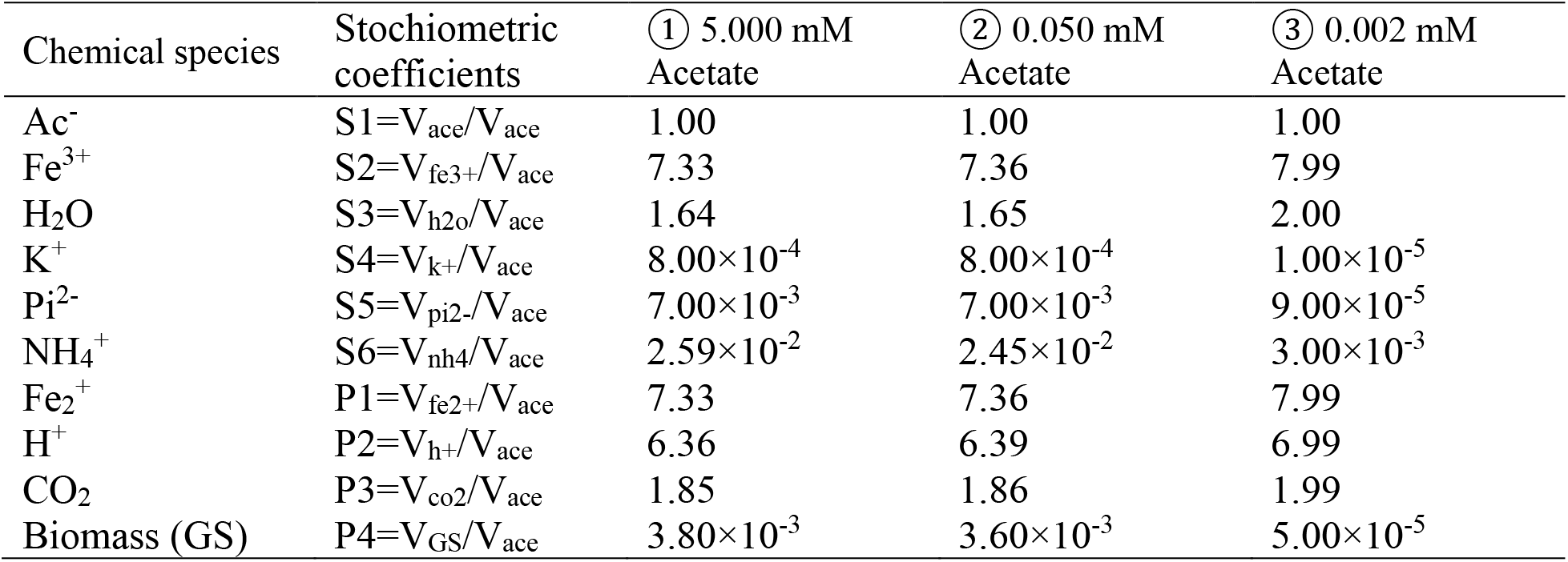
Microbial redox reaction stoichiometry under different acetate concentrations estimated from exchange fluxes through intracellular metabolism.

Therefore, our model predicts that biomass growth should be fastest in the outer biofilm strata, near the biofilm/liquid interface. Healthy, fast-growing layers have been previously observed in these positions for non-electroactive biofilms.^44, 45^ Evidence for similar preferential growth at outer layers is also presented in the literature for similar flow systems^11, 45^ has led authors to conclude that electron transfer is not significantly restricted away from the electrode surface. This could be due to biofilm compaction under flow, leading to better charge transfer. This also could be due to an increased active BMC in regions accessible by advection but not diffusion, resulting in an increased biofilm conductivity under flow, because it has been reported that the electrical conductivity of a GS biofilm is highly dependent on the local microbial activities.^46^ Notably, work in a static reactor concluded the opposite, that limitations to electron transfer away from the electrode surface were responsible for reduced activity at the biofilm strata located away from the electrode surface.^47^ Future development of our model may include investigating the question related to biofilm conductivity in order to weigh in more definitively on this topic.

From a theoretical standpoint, the ability of the GS GEM to dynamically change the reaction stoichiometry based on local conditions (e.g., acetate concentration) can explain why the conventional M-M kinetics, which use the constant stoichiometry in Eq. 1 to define the microbial oxidation of acetate, led to an overestimation of I^bioflm^ by 10% in nutrient-rich conditions (Figure 1). From an application perspective, iGEM-FRTM may be helpful for designing a novel bioelectrochemical system (BES) that precisely controls the local cellular metabolism and its surrounding environment. For example, based on real-time simulations to track all local environments, it will be possible that we can dynamically adjust the flow of nutrients to optimize the local biofilm structure as well as the tradeoff between energy versus biomass yield. Moreover, since GEM can monitor up to 56 exchange fluxes (GS_v3.xlsx in ESI), iGEM-FRTM may provide an opportunity to control biofilm acidification, nitrification, CO_2_ emission and fixation to maximize the BES performance. Future studies can also take advantage of iGEM to explore other aspects of intracellular metabolism (e.g. intercellular communications through exchange fluxes), especially because concentration gradients exist within the GS biofilm.

Looking ahead, prediction of the dynamic changes in the microbial redox reaction stoichiometry and the mass transport characteristics will help to reliably predict the scaleup performance of an emerging MECs (microbial electrolysis cells) technology via advances over existing techno-economic analysis using fixed redox reaction stoichiometry,^48^ which will place unrealistic pressures on new technologies that could cause their failure. However, our study shows that MEC whole-cell (e.g., GS) efficiency can deviate considerably from conventional assumptions by as much as 8% when considering microbial responses coupled to spatio-temporal dynamics. Therefore, our work can help assess the market-readiness of various MEC configurations by improving the predictability of scaled-up MEC costs and performance through more accurate estimation of the reaction stoichiometry.

#### (2) Bottom-up iGEM-FRTM can reproduce experimentally observed currents based on GS metabolic status and mass transport

The iGEM-FRTM uses the bottom-up approach to calculate I^biofilm^ by adding up the local I^cell^ in all finite-sized geometric shapes defined by the model mesh (e.g., triangles or squares) that are linked by local mass transfer. The goodness of fit between the measured and simulated I^biofilm^ depends on how the model correctly describes the GS metabolic status and mass transport in the GS biofilm.

Previous analysis of mass transport using flow reactors^11^ did not consider diffusion and advection as a parallel modes of mass transport. This led to values for D that were high compared to the present dynamically optimized iGEM-FRTM approach, which did account for both mass transport modes. As a result of the present approach, the lower values of D are more reasonable, considering that diffusion in biofilms is expected to be low due to frustrated movement through tortuous paths,^49^ diffusion barrier around cells,^50^ molecular adsorption^51^ and the presence of ion-biofilm coulombic interactions.^16^ It is worth noting that the low D we obtained can explain the reported observations in static diffusion-only reactors, where the static conditions limited diffusion through a GS biofilm, resulting in acetate molecules being excluded from all but the top 15% of the GS biofilm.^13^ Moreover, the reduced D helps to achieve a better goodness of fit between the observed and simulated data in continuous flow microfluidics, where advection occurs inside the GS biofilm. The iGEM-FRTM indicates that the Péclet numbers (ratio of advective transport rate to diffusive transport rate) were always greater than 1, revealing for the first time that advection inside the GS biofilm has a significant contribution to mass transport, at least for microfluidic BESs, which contrasts with previous work in flow systems. In each mesh element, the metabolic status of GS cells is represented by the GEM constrained by the local environment. To our knowledge, this is the first bottom-up modeling of the electrical power generation in electrochemically active biofilms to reproduce what has been experimentally measured and uncover the underlying mechanisms behind the given phenomena, such as the mass exchange between intracellular and extracellular compartments and the separate contribution to I^biofilm^ by acetate diffusion and advection.

Such a bottom-up view of the electrochemistry in a GS biofilm enhances our understanding of the GS biofilm environments, which have long been mysterious. Moreover, the iGEM-FRTM provides subtle details about the acetate, active biomass, and flow distribution over the biofilm in response to the culture conditions, which is hard to experimentally detect. The reader is directed to Figure S6 in the ESI for flow and concentration profiles through the modeled GS biofilm, which demonstrate that more of a loosely packed biofilm (high values of φ, k and D) can support faster advective flow than a compacted biofilm (low values of φ, k and D), resulting in different concentration gradients and flow patterns inside and outside the loose biofilm from the compact biofilm.

#### (3) Bottom up iGEM-FTRM can reproduce experimentally observed kinetic parameter KM based on fundamental cell-scale values

While it is a major step forward of the iGEM-FRTM to both reproduce experimental measurements of current (I^biofilm^) and to elucidate fundamental mechanisms involved, there remains the question of the origin of the order of magnitude difference between certain model parameters (e.g., K_M_) used at the biofilm-scale and the cell-scale. In addition, accurate reproduction of K_M_ parameter should demonstrate experimentally observed reductions as flow rate is increased. Therefore, the iGEM-FRTM was used to further identify a relationship between the M-M constant at the cell-scale 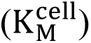 and at the biofilm-scale 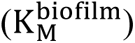 based on the simulated I^biofilm^ profiles.

The I^biofilm^ profiles were estimated as the sum of all local I^cell^ calculated with an experimentally determined 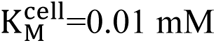 (a fundamental M-M constant for planktonic GS cells)^31^ for GEM. We then estimated 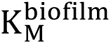 at each flow rate based on the M-M kinetics across a range of acetate concentrations (Figure 5A-5G), thus enabling comparison to values found experimentally 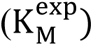.^21, 23^ As such, we found that the iGEM-FRTM simulations starting from 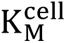 can follow the reported biofilm-scale M-M kinetics,^21, 23^ specifically generating 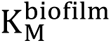 versus flow rate profiles that closely reproduced the observed exponential decreases in 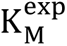 with increasing flow rate (often reported as apparent K_M_, or K_M(app)_). ^23^ A wide difference between the lower and higher bounds of 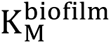 at low flow rates was found to be caused by an overestimation of I^biofilm^ under high acetate concentration (i.e., solid circles at 10 mM in Figure 5) due to omission of the biofilm acidification in the current version of the iGEM-FRTM. The next version of iGEM-FRTM that traces the fate of protons would narrow the difference between the upper-to-lower bounds. It should be noted that current literature points to the role of deacidification for the reduction in KM values with flow rate. While pH is not accounted for in this work, its connection to the charge-charge repulsion within the biofilm which we predict will swell the biofilm, were accounted for independantly through parameter optimizaiton in section discussed above.^34^

**Figure 5.**
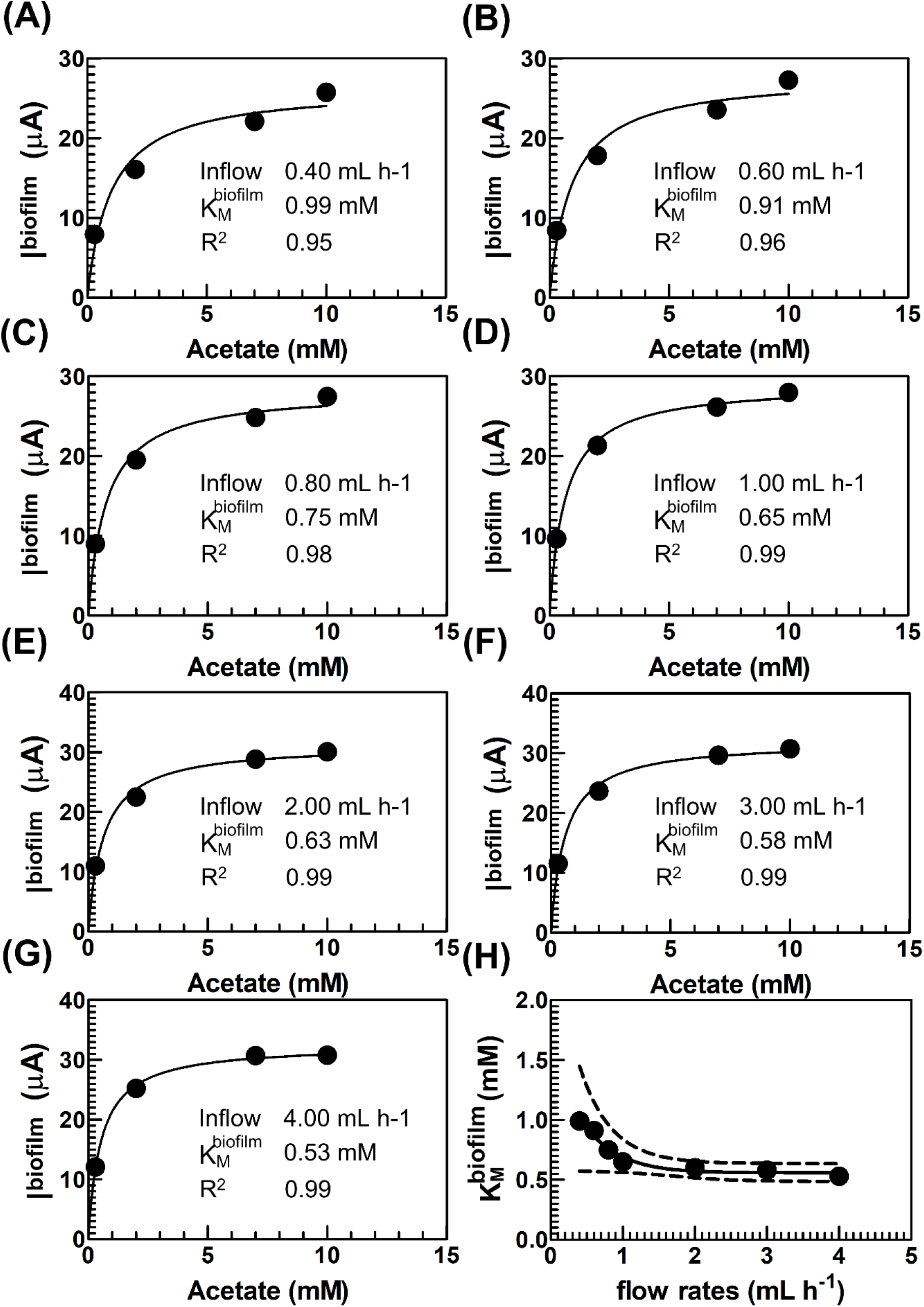
Relationship between 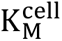 and 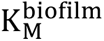. (A)–(G): estimation of 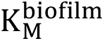 based on simulated electric intensity at steady state (I^biofilm^) across a wide range of inflow rates and acetate concentrations. I^biofilm^ values were predicted by adding up I^cell^ by the iGEM-FRTM with 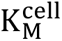 (0.01 mM) for GEM. (H): Change of 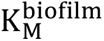 with flow rates. Solid circles: 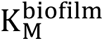 estimated from (A)–(G) at different flow rates, each across different acetate concentrations from 0.3 to 10 mM. Solid line: nonlinear regression of 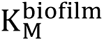 on flow rates. Dashed line: 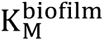 with one standard deviation of uncertainty.

Thus, the nature of 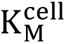 and the derived 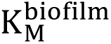 shows another advantage of the iGEM-FRTM as a predictive model for forecasting future biofilm behavior with the aid of the fundamental M-M constant and the existing biofilm status. This predictive function cannot be implemented by the conventional M-M kinetics to solve for condition-dependent parameters on a biofilm-scale by fitting the M-M equation against the measured I^exp^.

In addition to predictive modeling, given a specific biological process, GEM only requires the cell-scale M-M kinetics as model input, and their values are constant, independent of the biofilm conditions. Therefore, this approach can significantly reduce the inherent complexity for parameterizing the apparent kinetic parameters that are condition dependent at the biofilm-scale.

### Conclusion

In this study, we unified cell-scale metabolic activity with observed group behaviors in GS biofilms by integrating advanced modeling and experiments for a microfluidic system. The full microfluidic model couples mass transport with biological kinetics at the cell-scale. We used this model to capture the complex feedback between mass transport and metabolic status inside GS biofilms that are otherwise difficult to characterize beyond the cell-scale.

Despite the rise of flow-based BES devices, advection within GS biofilms has been largely neglected in the literature. Accounting for advection within GS biofilm in this work was critical in reproducing the experimental results, leading to the conclusion that in some previous studies which attributed to diffusion as the sole mass transport mechanism led to an overestimation of D. Moreover, we also predicted dynamic changes to biofilm structure and related mass transport based on changes to flow rate and nutrient concentration. We also provide insights into metabolic spatial variability, demonstrating the ability to map local metabolism and coulombic efficiency to different positions. Finally, we justified the magnitude differences in M-M constant K_M_ between different length scales, revealing the predictive function of the iGEM-FRTM based on cell-scale M-M constants to explore the unknown.

The unification of cell-scale metabolic activity and biofilm-scale behaviors identified the critical role played by local biofilm properties and metabolism. Such an in-depth understanding of the local biofilm environments may be helpful in formulating iterative model-based experimental optimization for biotransformation by engineering of the local biofilm structure or precise control of the cellular metabolism.

## Materials and methods

### Microfluidic device

The electrodes included a hand-cut graphite working electrode (WE) and counter-electrode (CE) (GraphiteStore.com Inc., USA) and a gold-coated polystyrene pseudo-reference electrode (RE) (fabricated via electroless deposition). After fabrication, all prepared electrodes had dimensions of 3 mm by 20 mm. We followed a report that demonstrated the method for embedding the electrodes at the bottom of a polydimethylsiloxane (PDMS) (Dow Corning, Canada) microchannel,^52^ which was defined by a photolithographic mold (FlowJEM Inc., Toronto, Canada) featuring a single channel with dimensions of 30 mm (length), 2 mm (width), and 0.4 mm (height). After embedding, the fluid-exposed electrode surface area for all electrodes was 3 mm by 2 mm, as shown in Figure 6. The gold pseudo-RE was placed upstream to ensure that the conditions were constant during the long-duration growth culture and bioelectrochemical experiments. The final device is the same as used in previous published works.^21, 25, 52^ Extensive tests were conducted previously into the stability of a microfluidic device with the same pseudo-RE and its positioning, which validated that the reference voltage was constant throughout the experiment (+400 mV vs. Ag/AgCl with 3 M KCl).^21, 25^ Cleaning and sterilization of the channel with a 70% ethanol solution and of the electrodes with 1 M HCl were performed before sealing.

**Figure 6.**
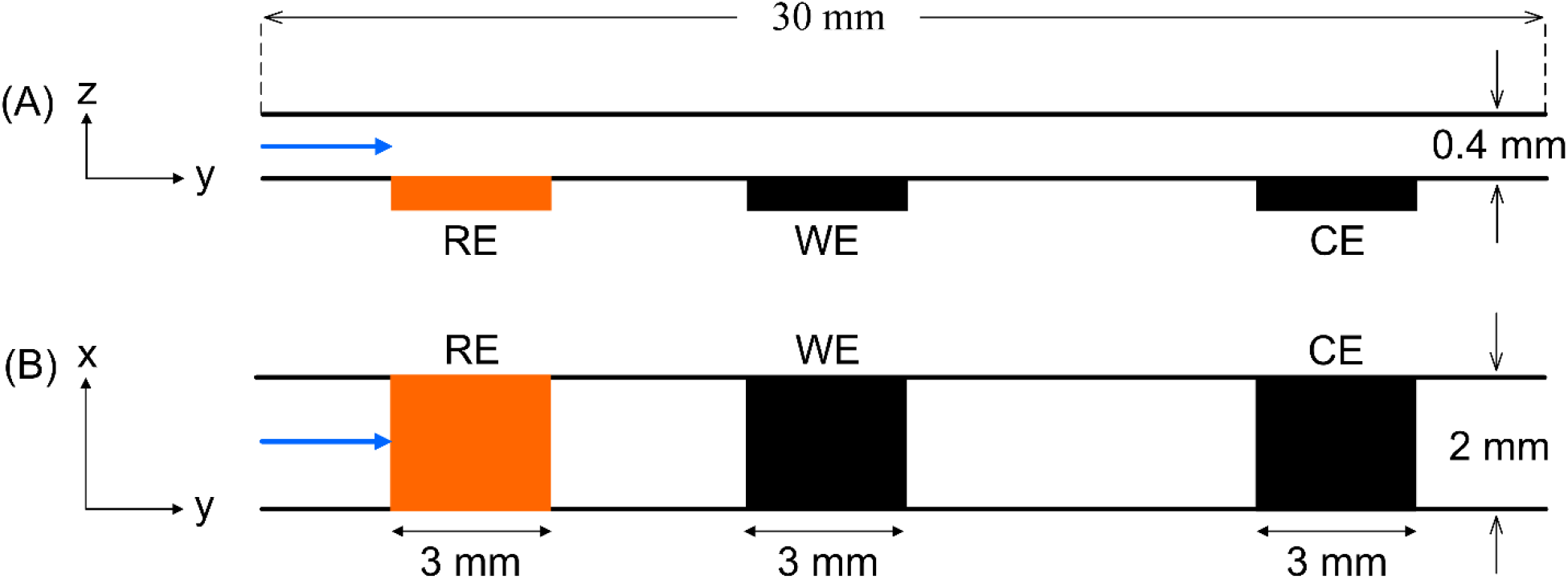
Schematic of a microfluidic channel from (A) the side view (z-y plane) and (B) the top view (x-y plane). Channel and electrode dimensions are given. Dimensions and electrode spacing are not to scale.

The microfluidic electrochemical cell was sealed with a microscope slide by exposure to air plasma (PCD-001Harrick Plasma, Ithaca, NY, USA). Because the microfluidic device was fabricated in PDMS, which is known to be porous to small molecules, including O_2_, an anaerobic enclosure (McIntosh and Filde’s, 28029 Sigma-Aldrich) was filled with anaerobic gas (20% CO_2_ and 80% N_2_) as done previously.^21, 25^ The device was placed in the enclosure, and sterile perfluoroalkoxy connective tubing (PFA tube 1/16, Hamilton Inc., Canada) and electrical connections were attached via airtight feedthroughs in the enclosure cap. To minimize air diffusion through the connective tubing outside of the enclosure, a layer of gas-impermeable tape (Loctite 249 Anaerobic Blue Threadlocker Tape, Medium Strength, Henkel Corp., Mississauga, Canada) was applied to the tubing, which was subsequently covered by epoxy glue. The electrical leads were fixed to the graphite WE and CE via alligator clips, and the gold RE was connected by solder with a protective coating of epoxy to physically stabilize the connection. The inlet tube was connected to a 50 mL glass syringe via connector assemblies (P-200x, P-658, IDEX, WA, USA). Syringe pumps (PHD 2000, Harvard Apparatus, Holliston, MA, USA) were used in controlled liquid injection. Before inoculation, the microfluidic channel and tubing were rinsed with sterile distilled water for 1 h at 1 mL h^-1^.

### Bacterial culture and device inoculation

Frozen samples of *Geobacter sulfurreducens* (strain PCA, ATCC 51573) were cultured under controlled temperature and deoxygenated conditions before injection into the microfluidic electrochemical device, as explained previously.^25^ The initial electrochemical growth of *G. sulfurreducens* was continuously monitored by a chronoamperometry electrochemical technique from 0 to 140 h (Figure S4). After the lag phase, a rapidly increasing anodic (oxidation) current was measured, indicating *G. sulfurreducens* biofilm growth at the WE. The WE was poised at an oxidative potential of 400 mV vs. Au (0 V vs. Ag/AgCl).

### Modeling approach for biological phenomena

We used three methods to describe the biological phenomena in the local environment. The microfluidics device used in this study was able to create a well-controlled, reproducible, 3-dimensional environment for a mature GS biofilm, with the metabolic status affected by limiting acetate concentrations. Hence, the biological phenomena in relation to the acetate oxidation kinetics were formulated in a 2-dimensional microfluidics model for simplicity but without loss of generality.

The first method used the M-M kinetics of acetate uptake to describe the overall GS growth and metabolism while treating the bacterial internal workings as a black box (Eq. 2 and Eq. 3).

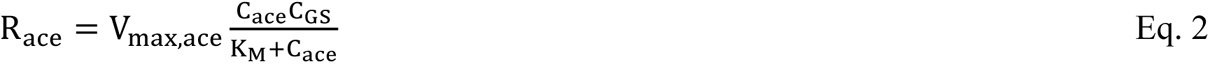

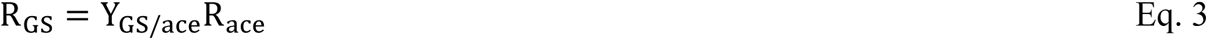

where R_ace_ and R_GS_ are the acetate uptake rates (mmol mm^-3^ h^-1^) and growth rates (g mm^-3^ h^-1^), respectively, and C_ace_ and C_GS_ are the acetate (mmol mm^-3^) and biomass concentration (g mm^-3^), respectively.

The second method was based on the genome-scale metabolic model (GEM) by applying flux balance analysis (FBA) to the GEM of GS. In this work, we made three modifications to a previously published GEM ^27^ to ensure that the model was mass balanced and charge balanced and devoid of obvious name errors in the gene–protein-reaction (GPR) system. First, any orphan metabolites and their associated reactions were removed from the model. Next, mass and charge balancing were performed by adding the metabolite information from the BiGG (Biochemical Genetic and Genomic)^53^ database. If information about a metabolite could not be found, the relevant formula and charge were derived by mass and charge balancing of the reactions with which they are associated. For cytochromes, due to the absence of concrete knowledge, the formula and charge of the metabolites that were most extensively annotated in the BiGG database were used. Finally, any erroneous names for the gene–protein-reaction (GPR) association were removed from its respective reaction, which was assigned an empty string for unknown GPR association instead of the reaction names.

FBA computes the vector of reaction fluxes (mmol g^-1^ h^-1^) that maximizes a cellular objective function (i.e., maximization of the specific growth rate, h^-1^), subject to mass balance constraints at steady-state for reaction network stoichiometry *S* and nutrient uptake rate constraints (Eq. 4). Formally, FBA is formulated as the following linear program.

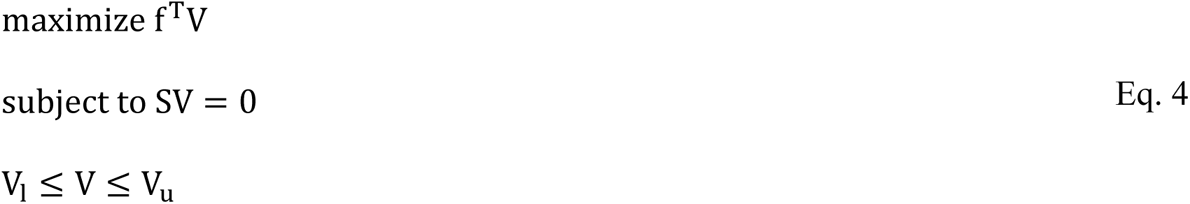

where V is the vector of n reaction fluxes (mmol g^-1^ h^-1^), f is the objective function vector, *S* is the stoichiometric matrix for *m* constraints, including the intracellular metabolite mass balance, and *l* and *u* are the lower and upper bounds on metabolic fluxes, respectively. For the acetate uptake reaction with metabolic flux V_ace_, we additionally constrained its flux via the M-M rate equation (Eq. 5). Thus, the acetate uptake flux (Eq. 6) and growth rate (Eq. 7) of GS cells are dependent on the local acetate concentration surrounding the CS cell surfaces, representing the local metabolic status.

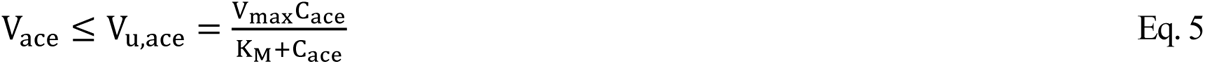

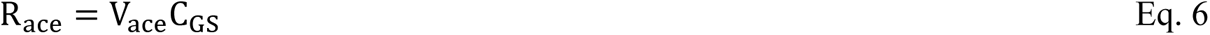

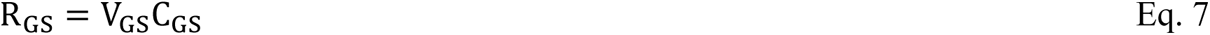

To avoid iteratively running the constraint-based GEM at each space grid point and at each time step, the third method based on the indirect implementation of FBA on GEM (iGEM) was used to reduce the computational cost due to numerous linear programming optimizations (LPs).

This approach used pre-calculated FBA predictions of the GS GEM model for the 10,000 possible concentrations of acetate ranging from 0.002 to 10 mM at a step size of 0.001 mM. These concentration points are environmentally relevant to those potentially observed in real microfluidics devices with respect to time and space. Next, we performed polynomial regression to obtain the relationship between the independent concentration variable (C_ace_) on a logarithmic scale and the dependent variable (electron transfer ratio and biomass yield) expressed as an nth degree polynomial in C_ace_. Here, the electron transfer ratio (ETR) is defined as the number of electrons (mmol) transferred to the anode per mmol of acetate oxidized, and the biomass yield (Y_bio/ace_) refers to the grams of dry-cell-weight biomass produced per mmol of acetate consumed. Thus, during execution of the microfluidics model, the polynomial function can be called at each time step and for each grid point to define ETR and Y_bio/ace_ at the given acetate concentrations. It should be mentioned that when acetate was the only growth limiting factor, the acetate uptake flux V_ace_(mmol g^-1^ h^-1^) computed by GEM, under concentrations of acetate from 0.002 mM to 10 mM, was found to be equal to its upper bounds (V_max_) as estimated by M-M kinetics. Hence, V_ace_ can be directly estimated by M-M, only with ETR and Y_bio/ace_ estimated by GEM. However, if the real uptake fluxes are different from their V_max_ due to other limiting factors, the above-mentioned polynomial regression method should be used to estimate the uptake fluxes.

### Modeling approach for flow and mass transport

In the full microfluidics model, we divided the microfluidics system into multiple virtual subdomains representing a three-electrode system with a reference electrode (RE), working electrode (WE), counter-electrode (CE), and the connection compartments between them (Figure 6). The GS biofilm was developed on the WE to an 80 µm thickness, with the headspace and other compartments filled with fluid. The GS biofilm was treated as partially permeable and porous media, with local pore spaces expressed as porosity (φ) and the ability for flow through the local environment expressed as permeability (k).

The model uses different equations to describe the flow and mass transport of different subdomains. The Navier-Stokes equation for computational fluid dynamics (CFD) and the diffusion-advection equation for mass transport are used for each discrete point in the free-flow subdomains (SD1, SD2, SD4, SD5 and SD6 in Figure 7). The Navier-Stokes-Brinkman equation for CFD (Eq. 8) and the diffusion-advection-reaction equation (Eq. 9 and Eq. 10) for reactive-transport are applied to each discrete point in the biofilm subdomain (SD3).

**Figure 7.**
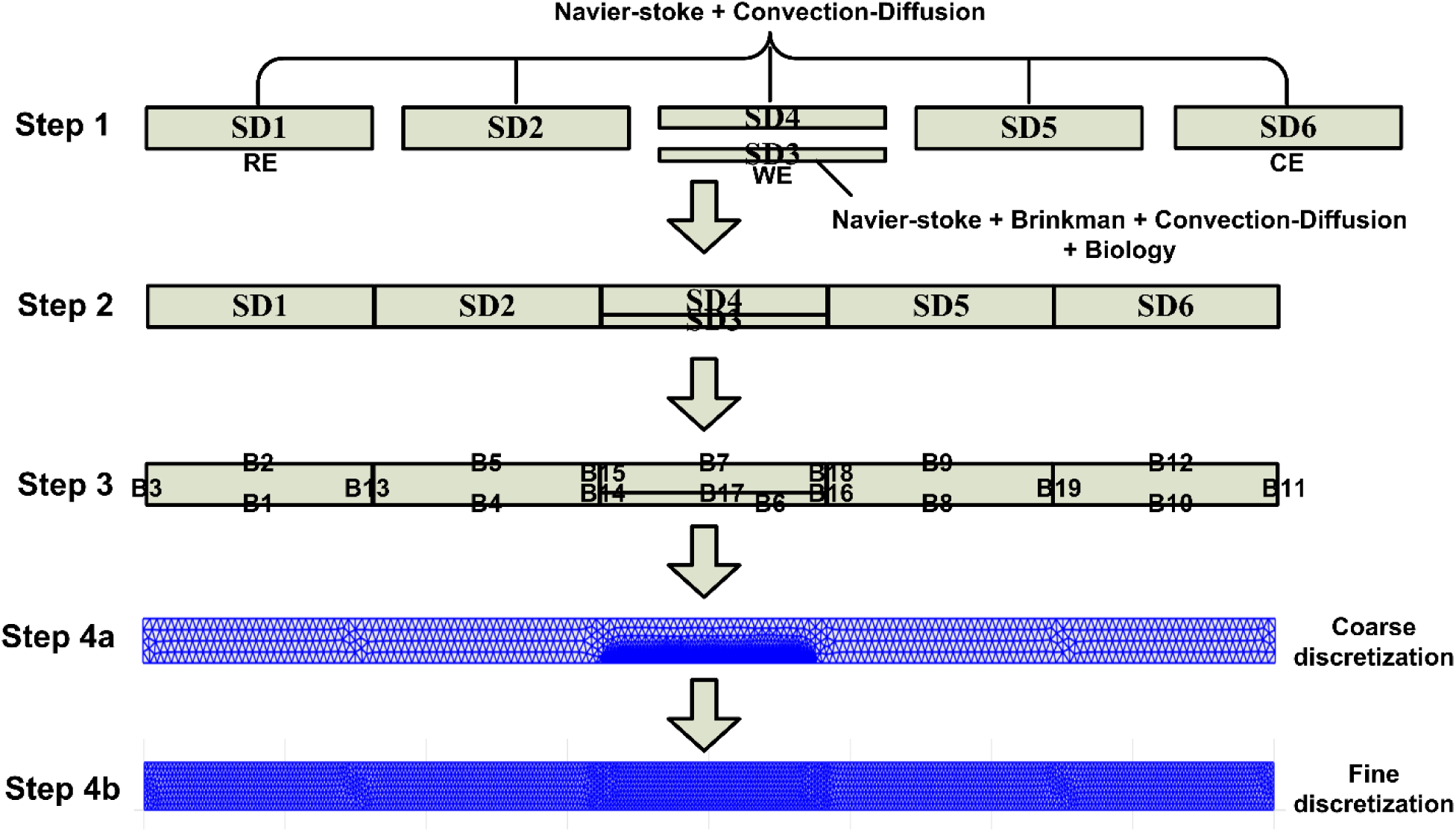
Microfluidics model designed to integrate GEM in local environments with flow and mass transport. SD1–SD6: subdomains representing the 3-electrode system with electrode (RE), working electrode (WE) and counter-electrode (CE). SD3: biofilm; SD4: headspace over the biofilm.

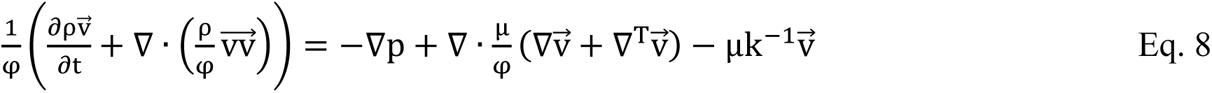

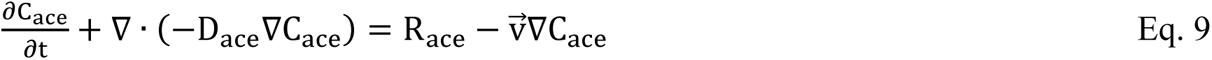

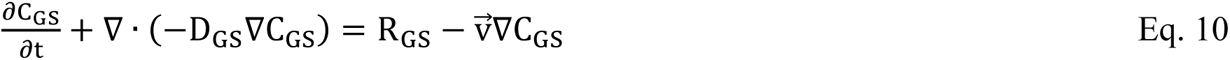

where φ is the porosity, 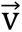 is the flow velocity vector, ρ is the fluid density, p is the static pressure, k is the permeability, D_ace_ and D_GS_ are the diffusion coefficients for acetate and biomass, respectively, μ is the dynamic viscosity, and R_ace_ and R_GS_ are local acetate uptake rates and growth rates for GS, respectively. All parameters describe the physical and biological phenomena in local environments, represented by the discrete points in the finite element method, and D_GS_is set to zero.

### Coupled physics and biological phenomena in microfluidics model

The full microfluidics model (GEM-FRTM) was built in FEATool Multiphysics (https://www.featool.com/) version 1.15 for Matlab. The block-coupling design by FEATool Multiphysics enables the model building pipeline (Figure 7) to assemble the virtually separate compartments into a full microfluidics simulator. These separate compartments are described by different governing equations for a three-electrode system connected by internal boundary conditions to satisfy the mass and flow continuity across the block interfaces. The Featool model for the full microfluidics device and the model structure readable by Matlab are provided in ESI (GS_microfluidics_model.fea and GS_microfluidics.mat).

The acetate uptake rates (R_ace_) and GS growth rates (R_GS_) computed by the M-M, GEM or iGEM models can be coupled to physical equations, as shown in Eq. 9 and Eq. 10. The finite element method (FEM) was used to solve the coupled mathematical equations based on space discretization generated by GRIDGEN2D (https://www.featool.com/doc/gridgen_8m). The FEM grid resolution and time step were optimized so that the model output did not change with the spatial temporal resolution.

The acetate concentrations, yield and transfer coefficients in space and time were predicted by the full microfluidics model, and the overall electricity production at steady state (i.e., simulated I^biofilm^), which can be compared to the measured current (I^exp^), was estimated for each flow condition as the sum of all simulated local current (I^cell^) based on the discrete grid points (Eq. 11).

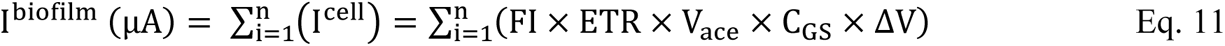

where n is the number of the discrete points, FI represents the flux to current constant, obtained by 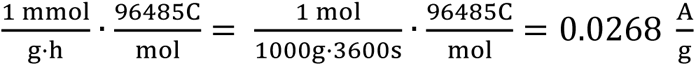, R_ace_ is the cell-scale acetate uptake rate (mmol mm^-3^ h^-1^), ΔV is the local volume for a discrete point, and ETR represents the electron transfer ratio from electron donor (acetate) to electron acceptor.

For simplicity without loss of generality, Fe^3+^ is chosen to be an extracellular electron acceptor in the GS GEM, with the same role as the outer-membrane cytochromes in the biofilm matrix to carry electrons towards the anode to generate current. The ratio of the Fe^3+^ reduction flux (mmol g^-1^ h^-1^) to acetate oxidation flux (mmol g^-1^ h^-1^) in the local environment represents ETR.

### Separate contributions of advection and diffusion to mass transport

The advection and diffusion fluxes in space and time were predicted by the full microfluidics model, and the overall advection and diffusion fluxes at steady state (i.e., 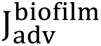 and 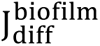 were estimated for each flow condition as the sum of all simulated advection and diffusion fluxes from individual local environment at the cell-scale (i.e., 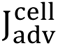 and 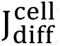 based on the discrete grid points (Eq. 12 and Eq. 13).

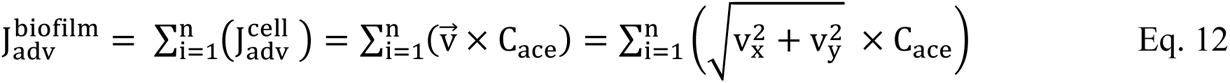

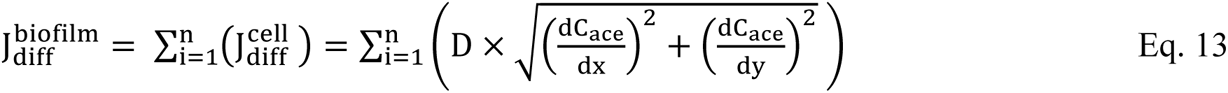

The Péclet number (P_eL_) was used to evaluate whether advection or diffusion has a significant contribution to the mass transport occurring in the biofilm (Eq. 14).

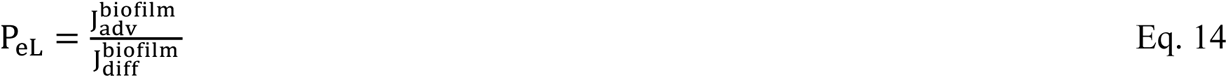

### Mass transport and biological kinetic factors

A global sensitivity analysis based on the Morris method was performed for 6 key local kinetic and mass transport input variables in the GEM-FRTM, including porosity (φ), diffusion coefficient (D), permeability (k), M-M parameters V_max_ and K_M_ and active biomass concentration (BMC). The detailed procedures of the Morris method for kinetic and mass transport input variables in FRTM are described elsewhere.^54^ In brief, the Morris sensitivity analysis was performed by sampling parameters within the ranges of possible values (Table 1) and calculating the individual and integrated effect of each parameter on the model outcome by Morris’ one-step-at-a-time (OAT) algorithm. The Morris analysis was performed by library “sensitivity” in R 4.1.0.

## Supporting information

ESI

## Nomenclature

BES: Bioelectrochemical systems
BMC: Local active biomass concentration (g dry cell weight per mm^-3^)
CFD: Computational fluid dynamics
D: Diffusion coefficient (mm^2^ h^-1^)
ETR: Electron transfer ratio (mmol of electrons transferred per mmol of acetate oxidized)
FBA: Metabolic flux balance analysis
FEM: Finite element method
FRTM: Flow and reactive transport model
GEM: Genome-scale metabolic model
GEM-FRTM: Genome-scale metabolic model coupled to flow and reactive transport model
iGEM-FRTM: Indirect genome-scale metabolic model coupled to flow and reactive transport model
GS: *Geobacter sulfurreducens*
iGEM: Indirect genome-scale metabolic model
I^biofilm^: Current produced in a GS biofilm at steady state (µA)
I^cell^: Current produced in a local environment around GS cell surfaces (cell-scale) in a GS biofilm at steady state (µA)
K: Local biofilm permeability (mm^-2^)
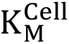: Michaelis-Menten constant at cell-scale (mmol mm^-3^) for GEM
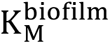: Simulated Michaelis-Menten constant at biofilm-scale (mmol mm^-3^)
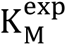: Measured Michaelis-Menten constant at biofilm-scale (mmol mm^-3^)
M-M: Michaelis-Menten kinetics model
V_max_: Maximum metabolic flux of the reaction (mmol g^-1^ dry cell weight h^-1^)
Y_bio/ace_: Biomass yield (g dry cell weight produced per mmol of acetate oxidized)
φ: Biofilm porosity (unitless)

## Supplementary legends

ESI: **Flow and reactive-transport modeling (FRTM-GEM)**

GS_v3.xlsx: **Modernized GS GEM in EXCEL format**

GS_v3.mat: **Modernized GS GEM in Matlab format**

GS_v3.xml: **Modernized GS GEM in Systems Biology Markup Language (SBML) format**

GS_microfluidics_model.fea: **Featool model for the full microfluidics device**

GS_microfluidics.mat: **Full microfluidics model structure readable by Matlab**

## Acknowledgements

The authors would like to thank Molly Gregas for the technical edits.

