## Supplementary material for "Unification of cell-scale metabolic activity with biofilm behavior by integration of advanced flow and reactive-transport modeling and microfluidic experiments": ESI

### ELECTRONIC SUPPLEMENTARY INFORMATION

#### ESI: Flow and reactive-transport modeling (FRTM-GEM)

##### (1) GS GEM Model (modernized version)

A modernized GS GEM that has updated GEM for GS (GS_v2) based on the original GS_v1.

| | |
|---|---|
| GS_v3.mat | 66 KB |
| GS_v3.xlsx | 55 KB |
| GS_v3.xml | 605 KB |

##### (2) Featool model for the full microfluidics device and model structure readable by Matlab

| | |
|---|---|
| GS_microfluidics_model.fea | 2,941 KB |
| GS_microfluidics.mat | 2,933 KB |

##### (3) Quality assessment of GS_v3 by MOMOTE [1]

***Table S1*** *Quality assessment of GS GEM model (GS_v3) by MOMOTE*

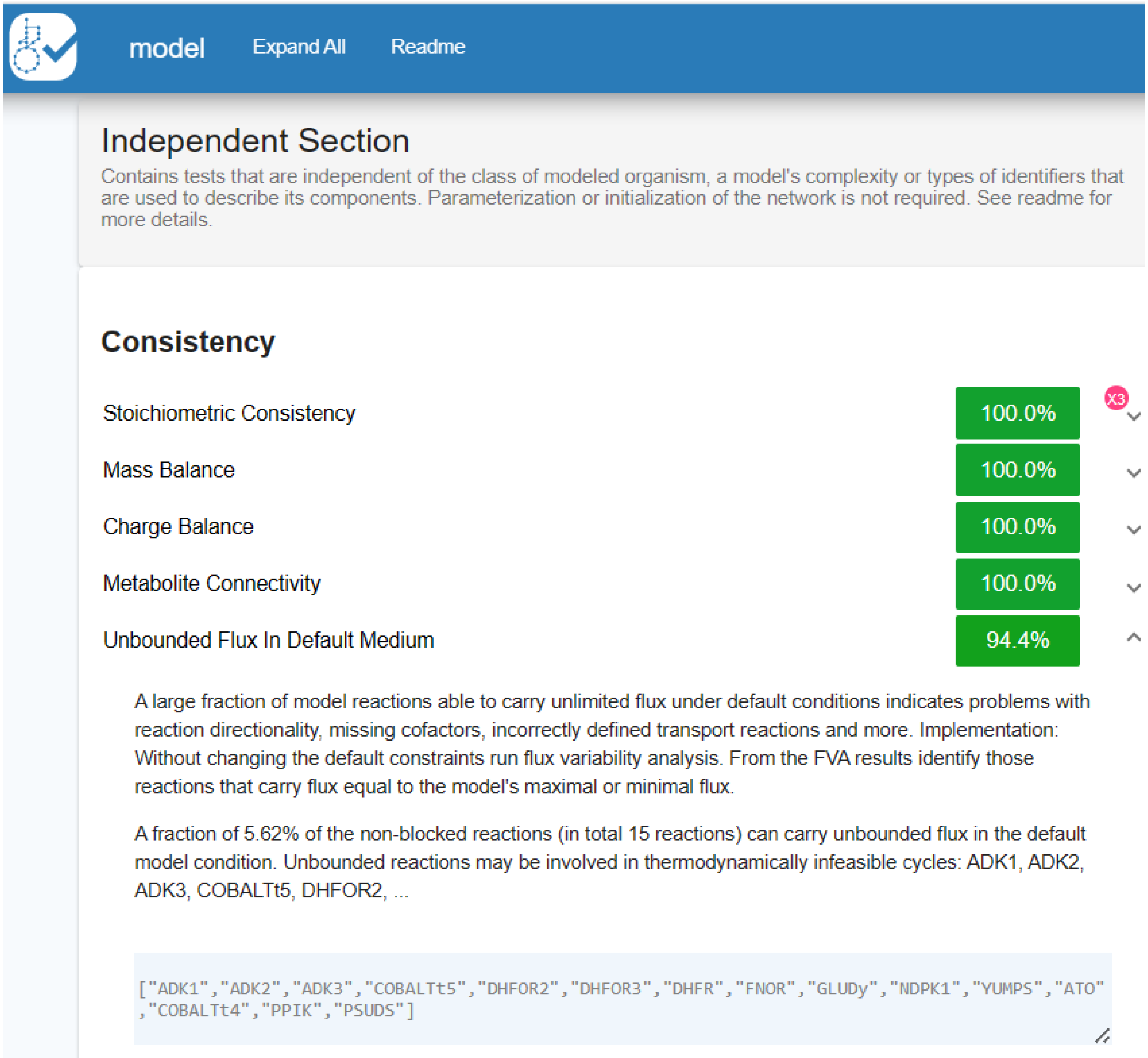

model
Expand All
Readme
Independent Section
Contains tests that are independent of the class of modeled organism, a model's complexity or types of identifiers that are used to describe its components. Parameterization or initialization of the network is not required. See readme for more details.
Consistency
Stoichiometric Consistency
100.0%
X3
Mass Balance
100.0%
Charge Balance
100.0%
Metabolite Connectivity
100.0%
Unbounded Flux In Default Medium
94.4%
A large fraction of model reactions able to carry unlimited flux under default conditions indicates problems with reaction directionality, missing cofactors, incorrectly defined transport reactions and more. Implementation: Without changing the default constraints run flux variability analysis. From the FVA results identify those reactions that carry flux equal to the model's maximal or minimal flux.
A fraction of 5.62% of the non-blocked reactions (in total 15 reactions) can carry unbounded flux in the default model condition. Unbounded reactions may be involved in thermodynamically infeasible cycles: ADK1, ADK2, ADK3, COBALTt5, DHFOR2, ...
["ADK1","ADK2","ADK3","COBALTt5","DHFOR2","DHFOR3","DHFR","FNOR","GLUDy","NDPK1","YUMPS","ATO","COBALTt4","PPIK","PSUDS"]

| | |
|---|---|
| SBML Level and Version | Level 3 Version 1 |
| FBC enabled | true |

The Flux Balance Constraints (FBC) Package extends SBML with structured and semantic descriptions for domain-specific model components such as flux bounds, multiple linear objective functions, gene-protein-reaction associations, metabolite chemical formulas, charge and related annotations which are relevant for parameterized GEMs and FBA models. The SBML and constraint-based modeling communities collaboratively develop this package and update it based on user input. Implementation: Parse the state of the FBC plugin from the SBML document.

The FBC package *is* used.

```
true
```

#### Basic Information

| | |
|---|---|
| Model Identifier | model |
| Total Metabolites | 466 |
| Total Reactions | 538 |
| Total Genes | 889 |
| Total Compartments | 2 |
| Metabolic Coverage | 0.61 |
| Uncoserved Metabolites | 0 |
| Minimal Inconsistent Net Stoichiometries | 0 |

#### (4) Loopless-FBA vs. LoopAllowed-FBA

MOMOTE indentified 5.62% (15 reactions) of the reactions that may carry unbounded fluxes, which may be involved in thermodynamically loops. The Loopless-FBA approach[2] by Mixed-Integer Linear Programming (MILP) Algorithm was used to elimiate these thermodynamically infeasible loops.

Pre-calculated Loopless-FBA predictions were done for the GS GEM model with the 10,000 possible concentrations of acetate ranging from 0.002 to 10 mM at a step size of 0.001 mM. Concentration-dependent ETR and $Y_{bio/ace}$ were obtained, as shown in Figure S1. For comparison, pre-calculated loop-allowed FBA predictions under 10,000 possible concentrations of acetate were shown in Figure 1.

Under test microfluidics conditions, it appears that the thermodynamically infeasible loops found in the GS GEM have no effect on ETR and $Y_{bio/ace}$, suggesting that both the loopless and loop-allowed FBA can be used in GEM-FRTM for this study. However, unphysical cycles in the metabolic network do need to be eliminated for energy balance, whose imbalance may be a risk of causing some mass and charge inequality for some GEM reactions. Unfortunately, the Loopless FBA approach is computationally expensive for large systems due to the Mixed-Integer Linear Programming (MILP) algorithm, which makes the coupling of GEM to FRTM impractical. The indirect GEM coupling method (I-GEM) proposed in this study provides a solution to addressing the challenge of such computational expenses.

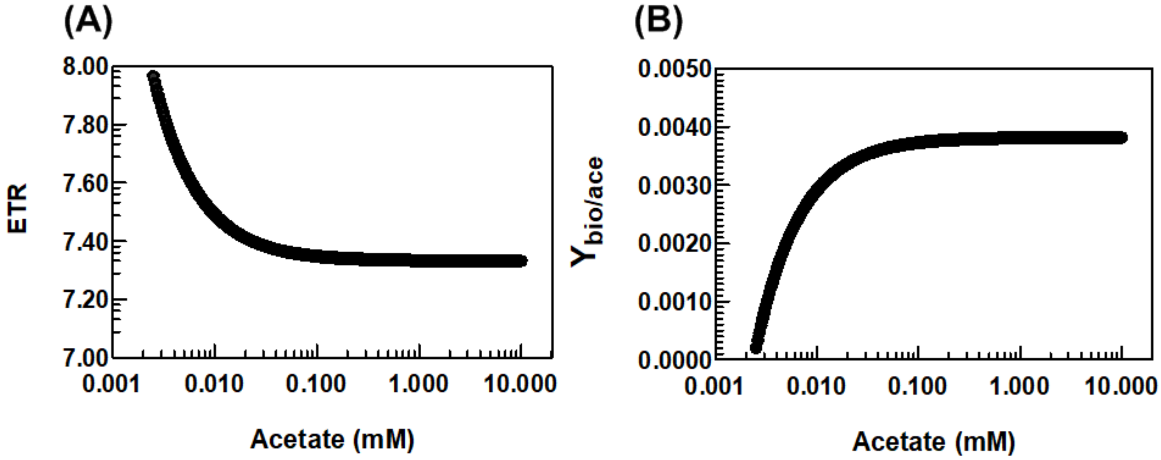


*Figure S1. Loopless FBA approach: (A) Semi-log regression of GEM-predicted electron transfer ratio (ETR, number of electrons transferred to current per oxidized acetate molecule) as a function of local acetate concentration; (B) semi-log regression of GEM-predicted biomass yield ($Y_{bio/ace}$, g dry-cell-weight biomass produced per mmol of acetate oxidized) as a function of local acetate concentration.*

#### (5) polynomial functions of electron transfer ratio (ETR) and biomass yield ($Y_{bio/ace}$) for the I-GEM method

$$ETR = 0.0020*(\log10(c))^6 + 0.0008*(\log10(c))^5 - 0.0013*(\log10(c))^4 - 0.0055*(\log10(c))^3 + 0.0060*(\log10(c))^2 - 0.0032*(\log10(c))^1 + 7.3340$$

$$Y_{bio/ace} = -0.000012*(\log10(c))^6 -0.0000049*(\log10(c))^5 +0.0000072*(\log10(c))^4+0.000032*(\log10(c))^3-0.000035*(\log10(c))^2+0.000018*(\log10(c))^1 +0.0038$$

#### (6) biomass estimation

**Volume of biofilm above the working electrode (WE):**

XYZ =3000 μm × 2000 μm × 80 μm = 4.8e+08 $\mu m^3$

**Volume of a bacterium (average):**

π × $r^2$ (radius of bacteria) × h (length of bacteria) =3.14 × $(0.25\ \mu m)^2$ × (1.7 μm) = 0.33 $\mu m^3$

**Number of total cells in the test biofilm:**

4.8e+08/0.33= 1.45e+09

**0.12–0.33 g dry weight per cubic centimeter of cell volume [3]**

average = 0.2250

**Dry cell weight per cell**

0.33*1e-09*0.2250/1e+03 = 7.4250e-14 g

**Total biomass weight**

7.4250e-14*1.45e+09

**Active biomass concentration (BMC)**

7.4250e-14*1.45e+09/(4.8e+08*1e-09) = 224e-06 g $mm^{-3}$

**BMC upper bound**

Adjust by average porosity 0.47: 0.53*224e-06 = 120e-06 g $mm^{-3}$

**BMC lower bound**

50% EPS and proteins: 120e-06*0.5 = 60e-06 g $mm^{-3}$

**BMC range**

[60e-06, 120e-06] g $mm^{-3}$ = [60, 120] g $L^{-1}$

#### (7) mass transport parameters for acetate

The diffusion coefficient (D) for acetate in the GS biofilm was initially set from 0.05 $mm^2\ h^{-1}$ to 0.10 $mm^2\ h^{-1}$, lower than the reported values in the literature (e.g. 0.7 ~ 9.0 $mm^2\ h^{-1}$).[4] Mathematically, mass transport in this study has been decoupled by advection and diffusion. A smaller D than the reported values ignoring the advection inside a biofilm is consistent with both advection and diffusion contributing to the mass transport of acetate.

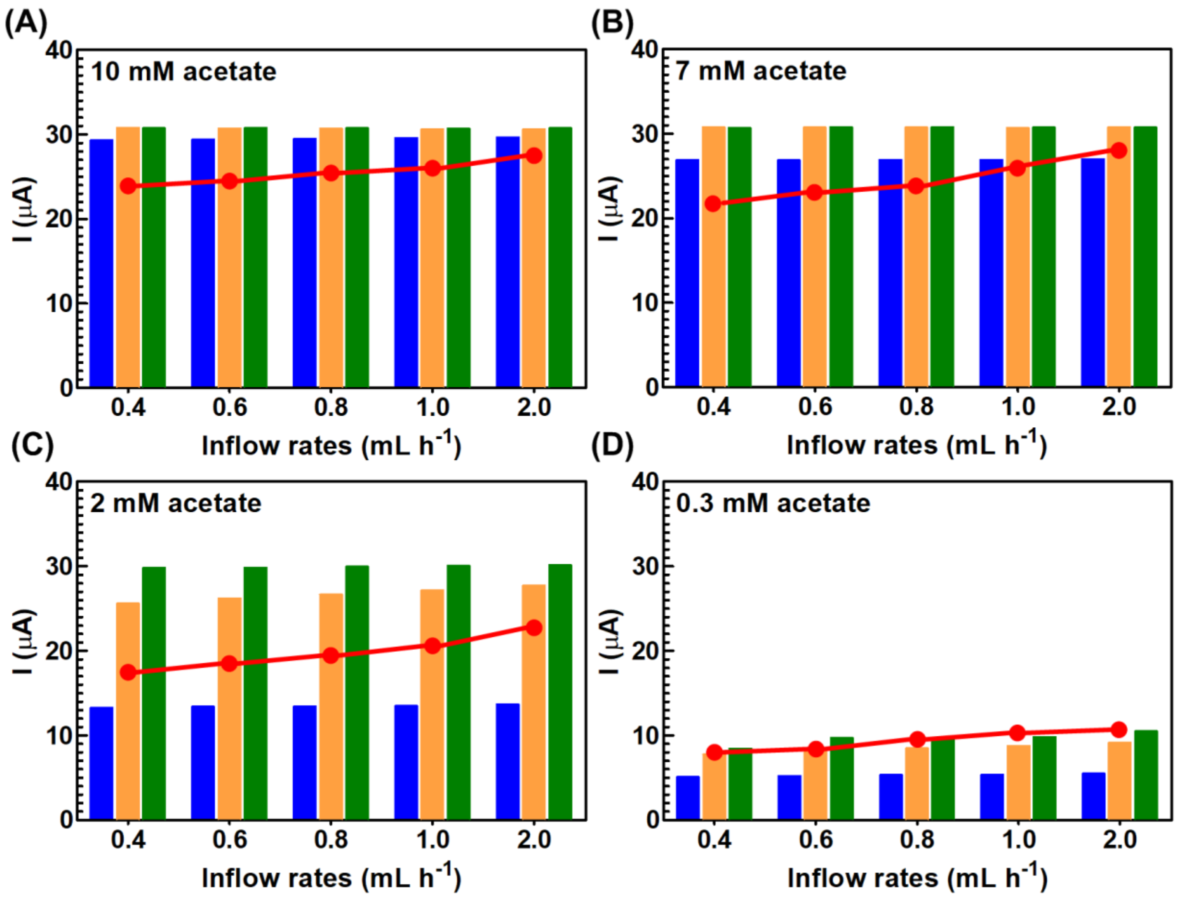


***Figure S2*** *The sensitivity of biofilm-scale currents simulated ($I^{biofilm}$, bars) and measured ($I^{exp}$, red dots) to inflow rates when diffusion is the only mass transport process inside the GS biofilm while flow only modulates the boundary layers around the biofilm when passing through other compartments. $I^{biofilm}$ simulated* at *local D:* ■ *0.05 mm$^2$ h$^{-1}$;* ■ *0.50 mm$^2$ h$^{-1}$;* ■ *5.0 (mm$^2$ h$^{-1}$);* ● *measured $I^{exp}$ from microfluidic experiments. Inflow acetate concentrations: (A) 10 mM and (B) 7.0 mM, (C) 2.0 mM, and (D) 0.3 mM.*

There is little information about the magnitude of k inside a bacterial biofilm. The k in a biofilm-porous medium system was reported to range from $10^{-9}$ to $10^{-1}$ mm$^2$.[5] Considering that the k values were estimated for porous medium systems, their magnitudes may not reflect the reality in the biofilm system alone. Moreover, the test biofilms (e.g., B. subtilis) are rich in extracellular biopolymers such as extracellular polymeric substances (EPS), DNA and proteins,[6] which are different from the GS biofilm characteristics. Thus, we first used our GEM-FRTM model simulations to preliminary determine the range of k from $2\times10^{-4}$ to $2\times10^{-3}$ mm$^2$ based on the experimental data of $I^{exp}$ across a wide range of inflow acetate rates and concentrations (see Figure 3 red band). This k range was consistent in magnitude with that used in a published FRTM for a general bacteria system.[7] Furthermore, the GEM-FRTM indicates that once k was further decreased beyond the lower bound at $2\times10^{-4}$ mm$^2$, the simulation cannot reproduce the stable $I^{exp}$ at 10 mM inflow acetate concentration after 0.07 hours but significantly decreasing

$I^{biofilm}$ to zero, suggesting the numerical issue due to the low input k values causing the incorrect mass transport performance for the GS biofilm system.

Bacterial biofilms are subject to reversible change in their structures with flow rates and substrate concentrations.[8-10] To reflect the loose structure of the GS biofilm under low flow rates and acetate concentrations, the D and k upper bounds were extended to 0.30 $mm^2\ h^{-1}$ and $2.5\times 10^{-3}\ mm^2$, respectively, to refine the mass transport parameters based on mechanical properties of the GS biofilm under these circumstances. We obtained a best match between $I^{biofilm}$ and $I^{exp}$ (Figure 3) by using the profiles of φ, D and k shown in Figure S3.

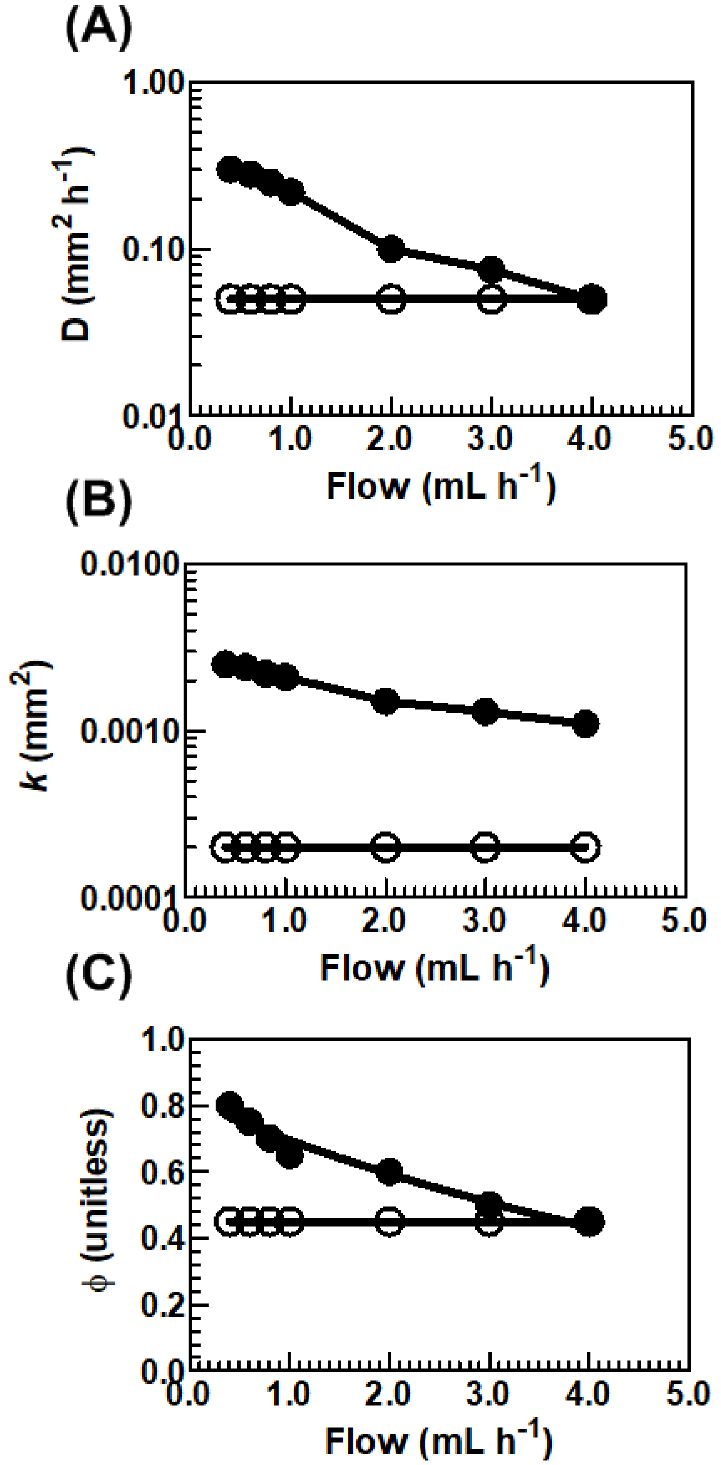


***Figure S3. (A)*** *The profiles of φ, D and k when a best match between* $I^{biofilm}$ *and* $I^{exp}$ *(Figure 3) was obtained.* ***Solid circle*** *(0.3 mM acetate);* ***open circle*** *(10 mM acetate). (A) diffusion vs. flow rates; (B) permeability vs. flow rates; (C) porosity vs. flow rates*

#### (8) details about homogeneous vs. heterogeneous biofilm

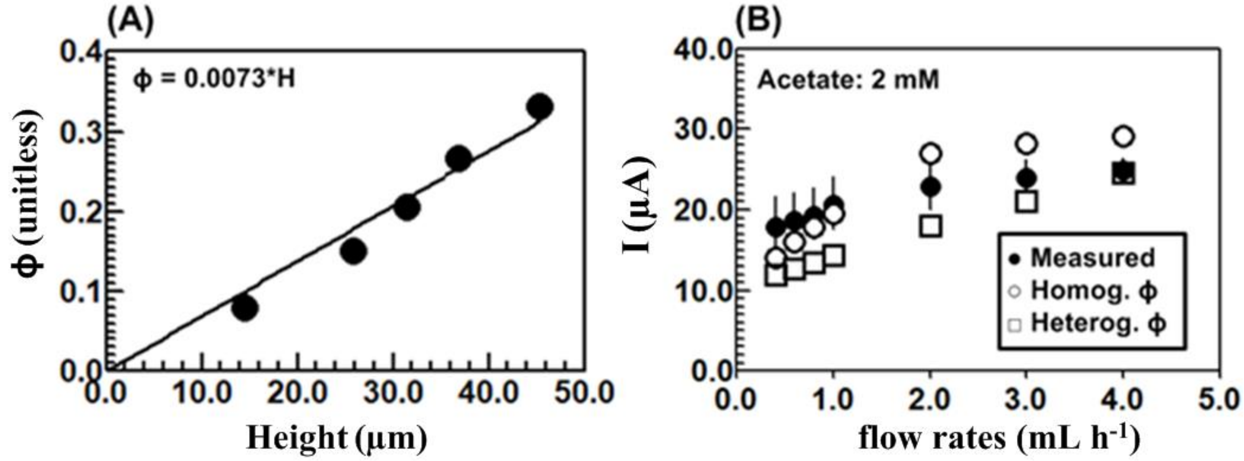


*Figure S4*. *Model extension: effect of biofilm structure heterogeneity (porosity) on the electricity production. (A) porosity represented as an empirical function of the biofilm height based on flow reactor experimental data;*[4] *(B) I (μA) vs flow rates under different biofilm structure*

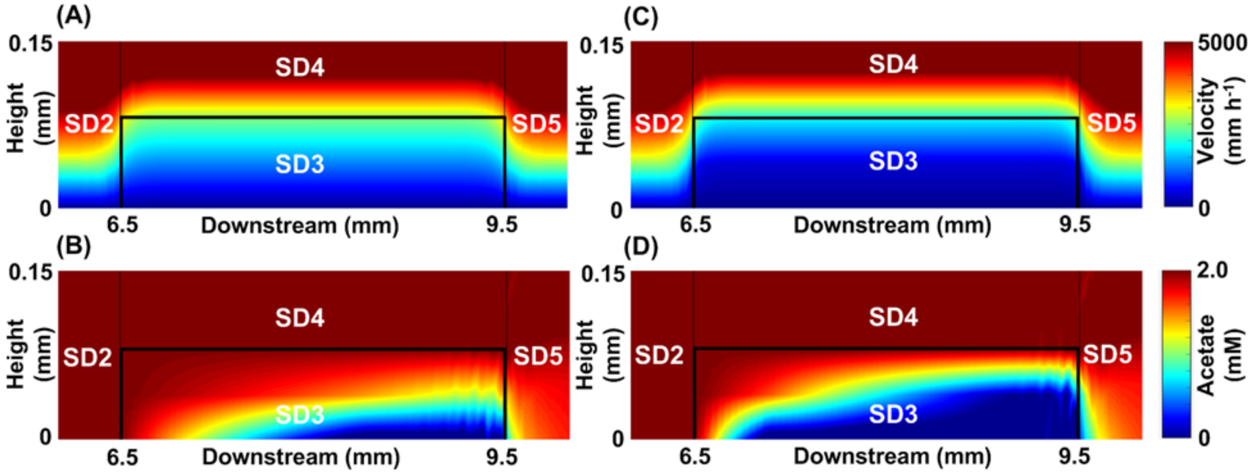


***Figure S5*. Homogeneous biofilm** *(constant φ) (A) flow field at flow rates of 4.0 mL $h^{-1}$ (B) acetate concentration map at flow concentration of 2.0 mM (C)* **Heterogeneous biofilm** *(φ as a function of a biofilm height; φ = 7.3*H(mm) based on experimental data reported for a ~ 400 μm thick GS biofilm): (C) flow field at flow rates of 4.0 mL $h^{-1}$ (D) acetate concentration map at flow concentration of 2.0 mM*

#### (9) details about loose vs. compact biofilm

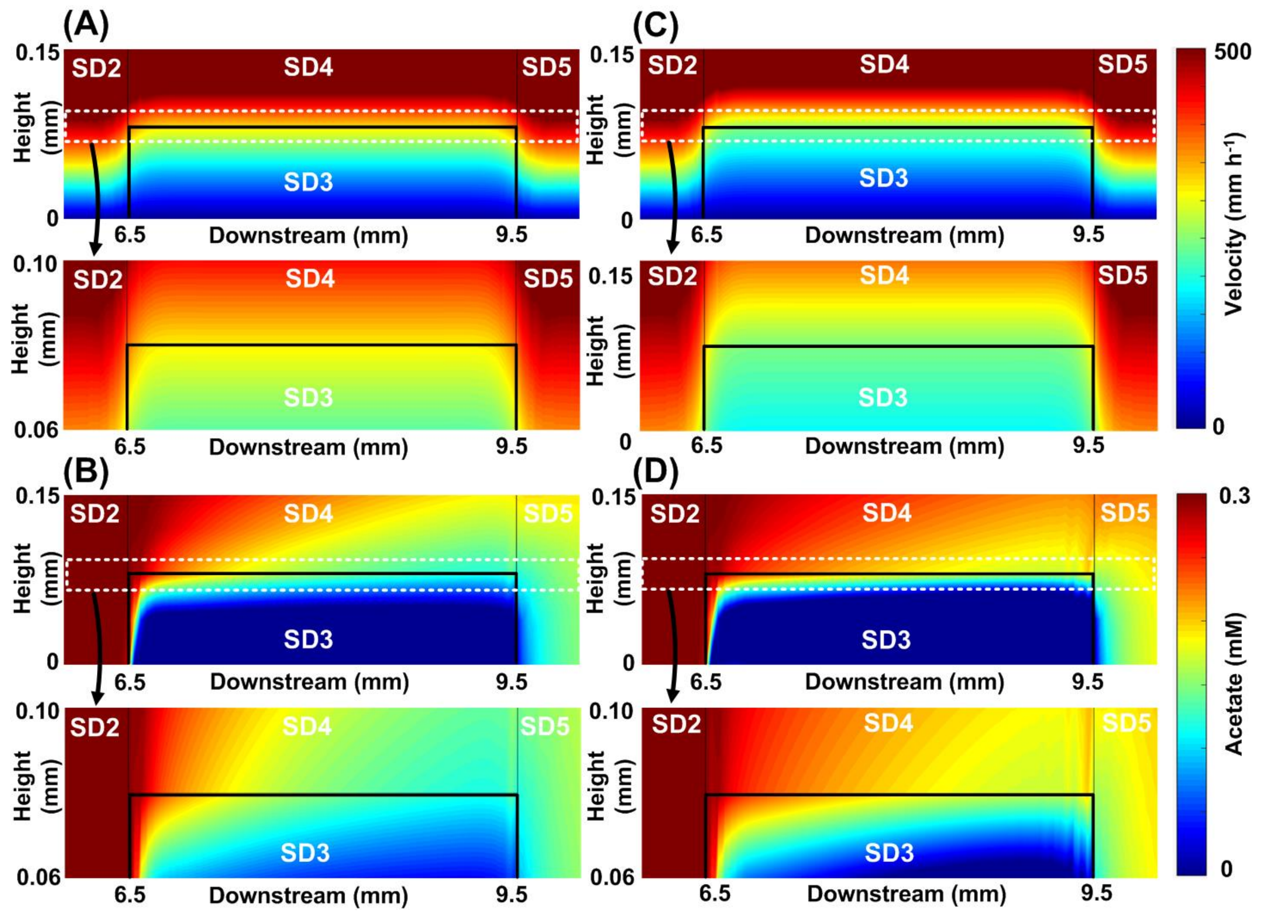


***Figure S6**. Different concentration gradients and flow patterns simulated inside and outside the loose biofilm from the compact biofilm.* **Loose biofilm** *(A) and (B) (D=0.30 mm$^2$ h$^{-1}$, φ=0.80): (A) top sub-figure (flow field from 0 to 150 µm height at inflow rates of 0.4 mL h$^{-1}$); bottom sub-figure: (zoomed view of the flow field from 60 to 100 µm height at flow rates of 0.4 mL h$^{-1}$) (B) top sub-figure (acetate concentration map from 0 to 150 µm height at flow concentration of 0.3 mM; bottom sub-figure (zoomed view of the acetate map from 60 to 100 µm height at flow concentration of 0.3 mM)* **Compact biofilm** *(C) and (D) (D=0.05 mm$^2$ h$^{-1}$, φ=0.45): (C) same as (A); (D) same as (B). SD2 and SD5: biofilm-free compartments; SD3: biofilm; SD4: headspace over the biofilm. See full subdomains (SD) in Figure 6.*

#### (10) Nature of mass transport D

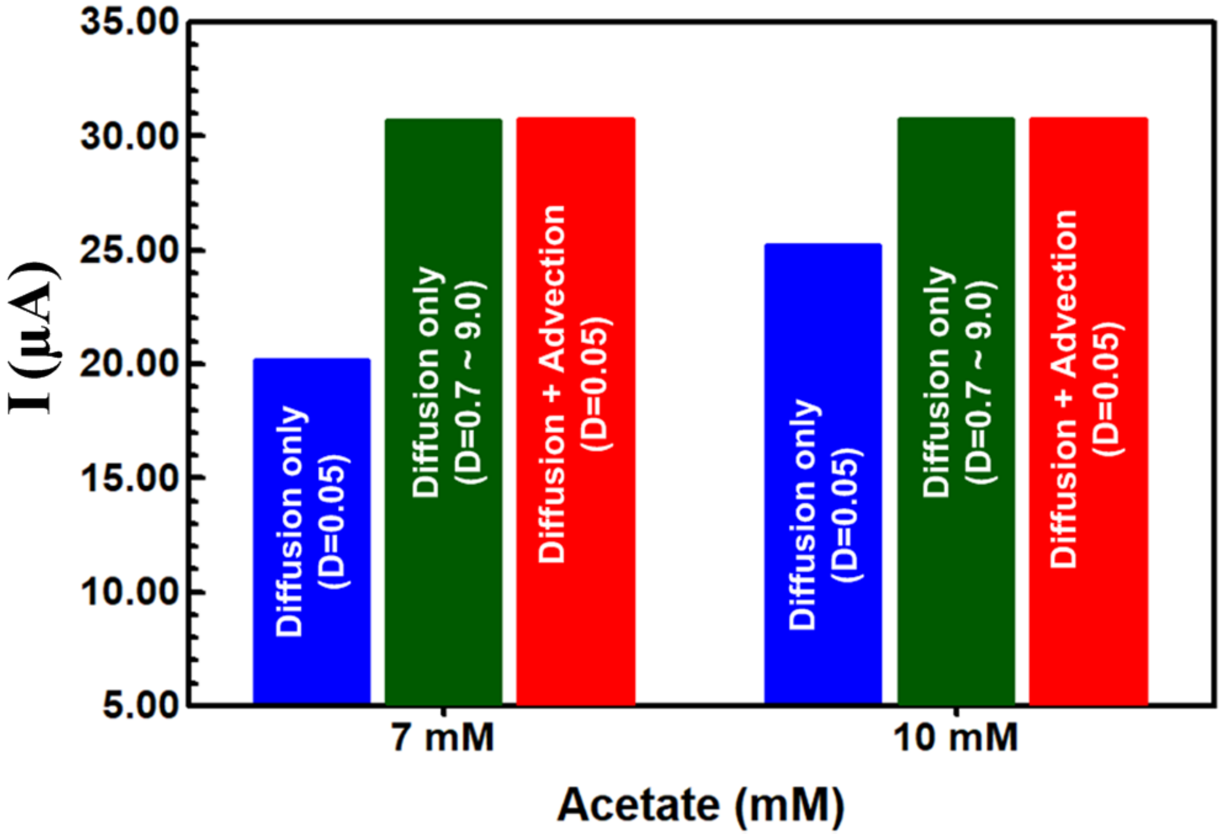


***Figure S7.*** *$I^{biofilm}$ estimated by GEM-FRTM with diffusion-driven-only and diffusion-advection simulations under reported experimental conditions.* [4] *Under reported experimental conditions (higher acetate concentration at 1 mL $h^{-1}$ of flow) where advection was omitted inside the biofilm, the diffusion-driven-only D has to be increased in GEM-FRTM to reach the $I^{biofilm}$ level obtained by both diffusion and advection contribution.*
